# A turquoise fluorescence lifetime-based biosensor for quantitative imaging of intracellular calcium

**DOI:** 10.1101/2021.06.21.449214

**Authors:** Franka H. van der Linden, Eike K. Mahlandt, Janine J.G. Arts, Joep Beumer, Jens Puschhof, Saskia M.A. de Man, Anna O. Chertkova, Bas Ponsioen, Hans Clevers, Jaap D. van Buul, Marten Postma, Theodorus W.J. Gadella, Joachim Goedhart

**Affiliations:** Swammerdam Institute for Life Sciences, Section of Molecular Cytology, van Leeuwenhoek Centre for Advanced Microscopy, University of Amsterdam, Amsterdam, the Netherlands; Department of Molecular Hematology at Sanquin Research and Landsteiner Laboratory, Academic Medical Centre, University of Amsterdam, the Netherlands; Oncode Institute, Hubrecht Institute, Royal Netherlands Academy of Arts and Sciences and University Medical Center, Utrecht, the Netherlands; Oncode Institute, Center for Molecular Medicine, University Medical Centre Utrecht, Utrecht, the Netherlands

## Abstract

The most successful genetically encoded calcium indicators (GECIs) employ an intensity or intensiometric readout. Despite a large calcium-dependent change in fluorescence intensity, the quantification of calcium concentrations with GECIs is problematic, which is further complicated by the sensitivity of all GECIs to changes in the pH in the biological range. Here, we report on a novel sensing strategy in which a conformational change directly modifies the fluorescence quantum yield and fluorescence lifetime of a circular permutated turquoise fluorescent protein. The fluorescence lifetime is an absolute parameter that enables straightforward quantification, eliminating intensity-related artifacts. A new engineering strategy that optimizes lifetime contrast led to a biosensor that shows a 3-fold change in the calcium-dependent quantum yield and a fluorescence lifetime change of 1.3 ns. Additionally, the response of the calcium sensor is insensitive to pH between 6.2-9. As a result, the turquoise GECI enables robust measurements of intracellular calcium concentrations by fluorescence lifetime imaging. We demonstrate quantitative imaging of calcium concentration with the turquoise GECI in single endothelial cells and human-derived organoids.

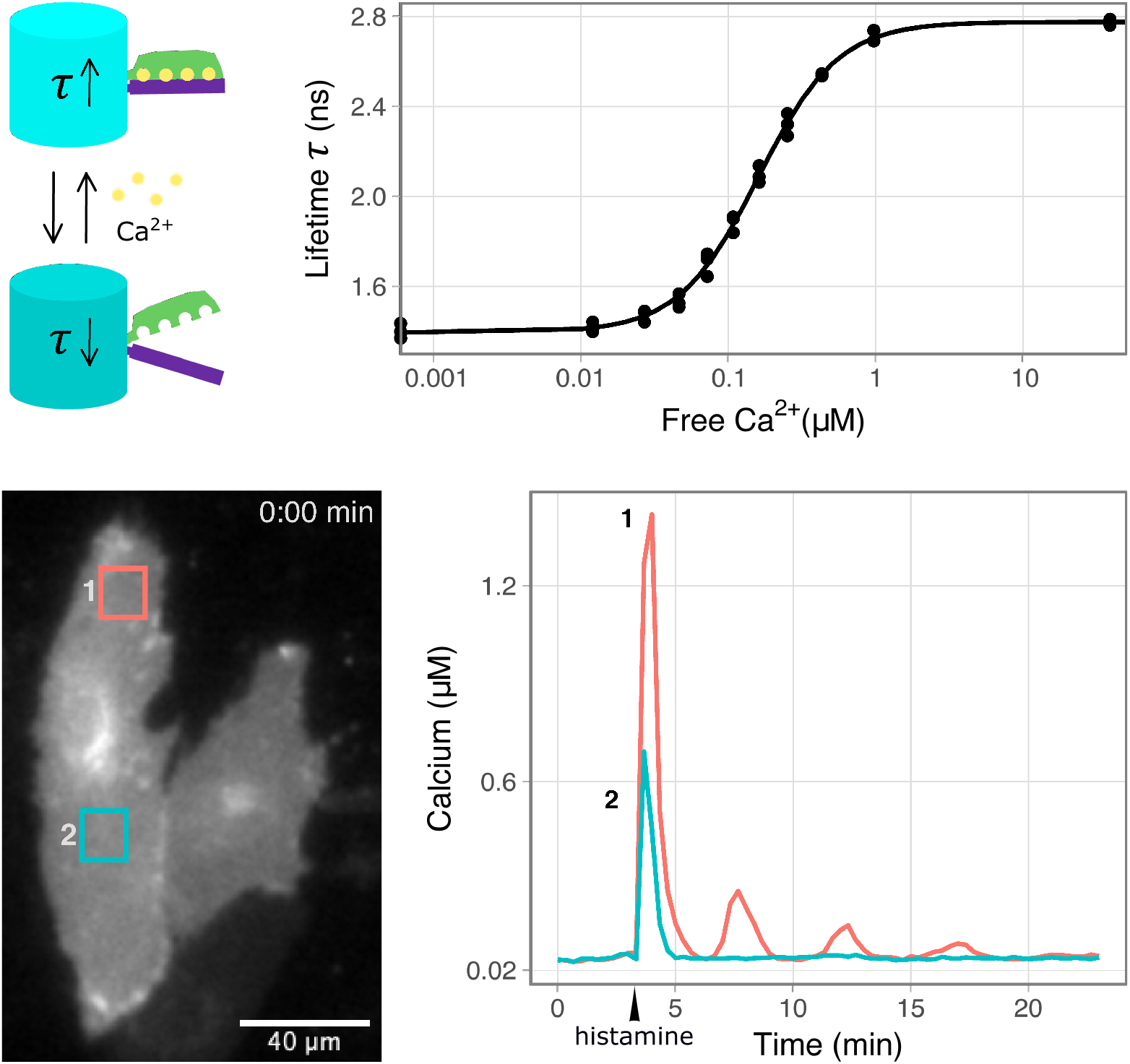

## Main text

Genetically encoded calcium indicators (GECIs) are popular tools for probing intracellular calcium levels^1^. Fierce engineering efforts have led to calcium probes with an impressive intensity contrast, enabling functional imaging in complex tissue and animals^2,3^. The probe design that delivers the largest intensity contrast uses a circular permutated green fluorescent protein (cpGFP) flanked by calmodulin (CaM) and a peptide that binds calcium-bound calmodulin^3–5^. The calcium-dependent interaction between calmodulin and the peptide results in a conformation change that is converted into a change in fluorescence intensity of the cpGFP. These probes are also known as GCaMP or GECO^6^. Alternatively, chemical dyes have been used for calcium sensing, but they are limited in application to cellular cultures and they are, unlike GECIs, not targetable to specific organelles^7^.

Despite their success, the GCaMP-type probes have limitations. First, the intensity-based read-out hinders quantification. The calcium levels modify the fluorescent intensity of the cpGFP. However, fluorescence intensity can also be changed by many additional factors, including photobleaching, sample movement, and changes in (local) probe concentration^8^, making it inherently difficult to quantify. Second, GCaMP type probes are sensitive to pH^9,10^. The binding of calcium changes the p*K*_*a*_ of the cpGFP, which results in a change in the protonation of the chromophore, which ultimately leads to the intensity change^11^. The p*K*_*a*_ of GECIs is near the physiological pH and therefore a change in intracellular pH also changes the intensity. Together, these factors complicate true quantitative imaging of calcium.

Efforts have been made to correct for some these effects by ratio imaging using a second fluorescent protein (FP), usually an orange or red variant^12,13^. Unfortunately, the ratio depends on the intensity of the second FP, which is turn is determined by its maturation^12^ (**Figure S1**). In addition, these sensors have inherited the pH sensitivity of their parents^12,13^. FRET-based probes are another alternative but have the same issues with unequal maturation rates of the two fluorescent proteins. Also, the widely used yellow FP as acceptor is relatively pH sensitive^14^.

We observed for both types of ratiometric sensors a high variability in emission ratio between cells (**Figure S1, Table S1**). This variability hinders quantification and demands establishment of the dynamic range for each individual cell. In addition, intensity and therefore ratio-imaging is dependent on instrumentation and therefore data obtained on different microscopes are not comparable. Lastly, FRET and ratio-imaging requires a large portion of the visible spectrum, reducing possibilities for multiplexing.

**Figure S1.**
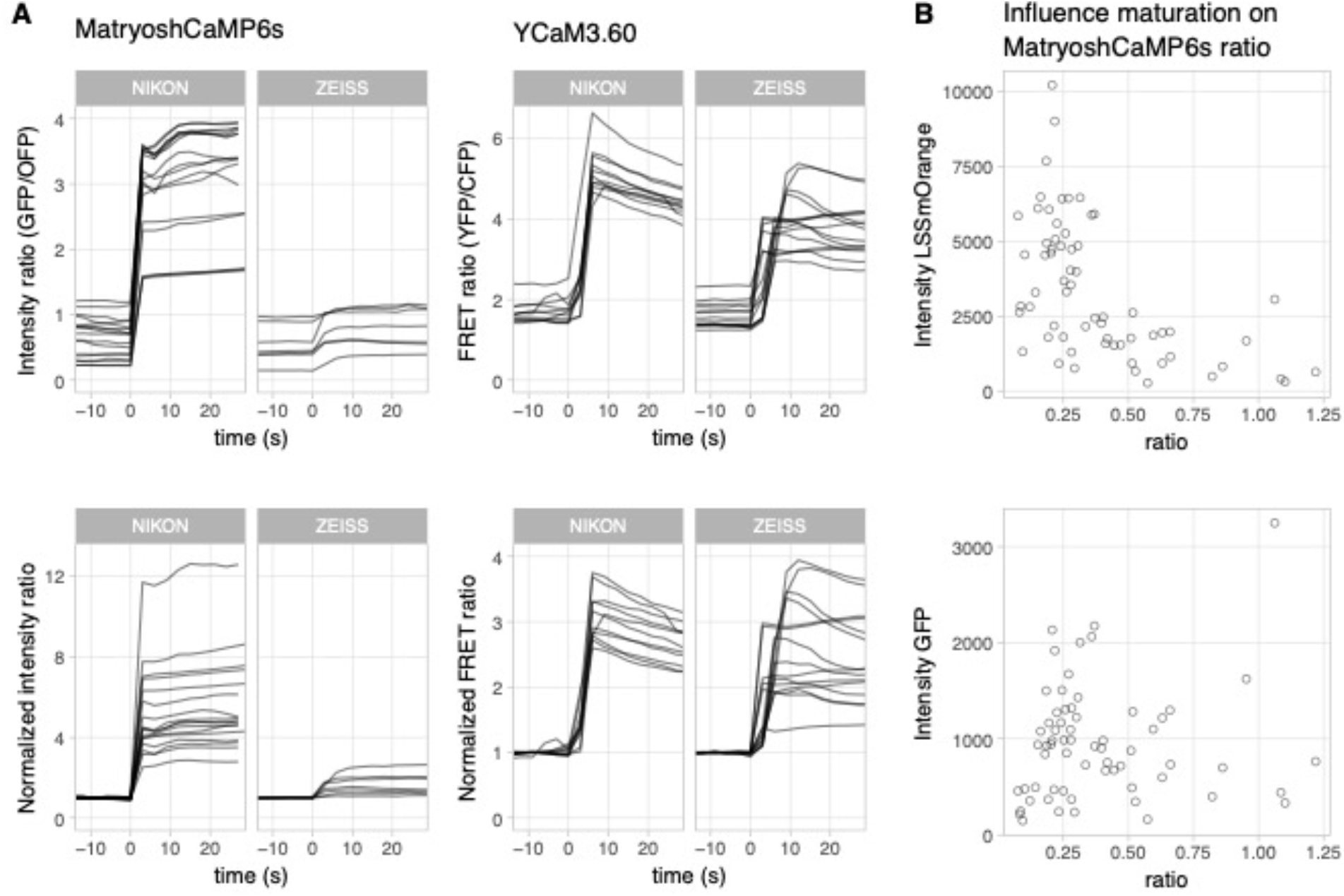
Ratiometric measurements of MatryoshCaMP6s and YCaM3.60 in HeLa cells. **A)** Timeseries of individual cells stimulated at t=0 with 14 mM ionomycin combined with 5 mM CaCl_2_ (*n*=7 to 17), taken on two different microscopes, a NIKON and a ZEISS setup. Top panels show the intensity ratio, bottom panels show the ratio normalized to the first three frames. Both ratio and normalized ratio give a highly variable output. **B)** The intensity of LSSmOrange of MatryoshCaMP6s (top panel) shows a clear influence on the measured ratio, which is not the case for the green fluorescence (bottom panel). Dots indicate individual cells (n=64).

**Table S1.** Variation of the output of different calcium sensors in HeLa cells. The respective fluorescent property was measured for each sensor before (pre) and after (post) addition of 14 mM ionomycin combined with 5 mM CaCl_2_. For lifetime measurements both the phase and modulation lifetimes are indicated. The mean, standard deviation (sd) and coefficient of variation (CV, sd divided by mean) of individual cells are given. CV is a number for the variation, independent of the absolute value, and can therefore be used to compare the variation of different types of readouts. Tq-Ca-FLITS published here was added for comparison and shows the lowest CV of all sensors.

| Sensor | Measured property | state | N | mean | sd | CV |
| --- | --- | --- | --- | --- | --- | --- |
| Tq-Ca-FLITS | phase lifetime (ns) | pre | 110 | 1.45 | 0.036 | 0.025 |
|  |  | post | 110 | 2.77 | 0.035 | 0.013 |
|  | modulation lifetime (ns) | pre | 110 | 1.84 | 0.055 | 0.030 |
|  |  | post | 110 | 2.94 | 0.034 | 0.012 |
| jRCaMP1b | phase lifetime (ns) | pre | 29 | 1.60 | 0.120 | 0.075 |
|  |  | post | 56 | 2.78 | 0.040 | 0.014 |
|  | modulation lifetime (ns) | pre | 29 | 2.35 | 0.103 | 0.044 |
|  |  | post | 56 | 3.02 | 0.055 | 0.018 |
| RCaMP1h | phase lifetime (ns) | pre | 31 | 1.05 | 0.049 | 0.047 |
|  |  | post | 69 | 2.74 | 0.040 | 0.014 |
|  | modulation lifetime (ns) | pre | 31 | 1.72 | 0.079 | 0.046 |
|  |  | post | 69 | 2.97 | 0.049 | 0.017 |
| MatryoshCaMP6s | intensity ratio (GFP/OFP) | pre | 7 | 0.54 | 0.291 | 0.536 |
|  |  | post | 7 | 0.83 | 0.307 | 0.368 |
| YCaM3.60 | FRET ratio (YFP/CFP) | pre | 15 | 1.58 | 0.305 | 0.192 |
|  |  | post | 15 | 4.08 | 0.720 | 0.177 |

The excited state fluorescence lifetime of a fluorescent molecule is generally not influenced by intensity-related factors^15–17^, and has been used successfully for quantitative imaging^18–22^. Consequently, GECI probes based on lifetime contrast would enable true quantitative calcium imaging, independent of equipment. However, most current GECIs, including GCaMPs and FRET-based GECIs, show hardly any or no QY or lifetime contrast (**Figure S2, Table S2**), despite a clear intensity change^23,24^. For the FRET-sensors this is caused by a close to 100% efficiency in energy transfer in one of the states of the sensor. In cpGFP based sensors the intensity change is predominantly caused by a change in extinction coefficient (*ε*), without a significant change in quantum yield (*QY*)^3,23^ (**Table S2**). This leads to an absence of lifetime change since the lifetime of fluorescent proteins is proportional to the *QY*.

Exceptions are RCaMP1h and jRCaMP1b, which are intensity-based calcium sensors that employ a red fluorescent protein instead of a GFP and they show a *QY* and lifetime contrast^25,26^. However, these probes display pH sensitivity in the biological range and a lower calcium sensitivity compared to other sensors. In addition, they have a very low intensity in the calcium-free state, complicating lifetime measurements (**Figure S3, Table S2**).

**Figure S2.**
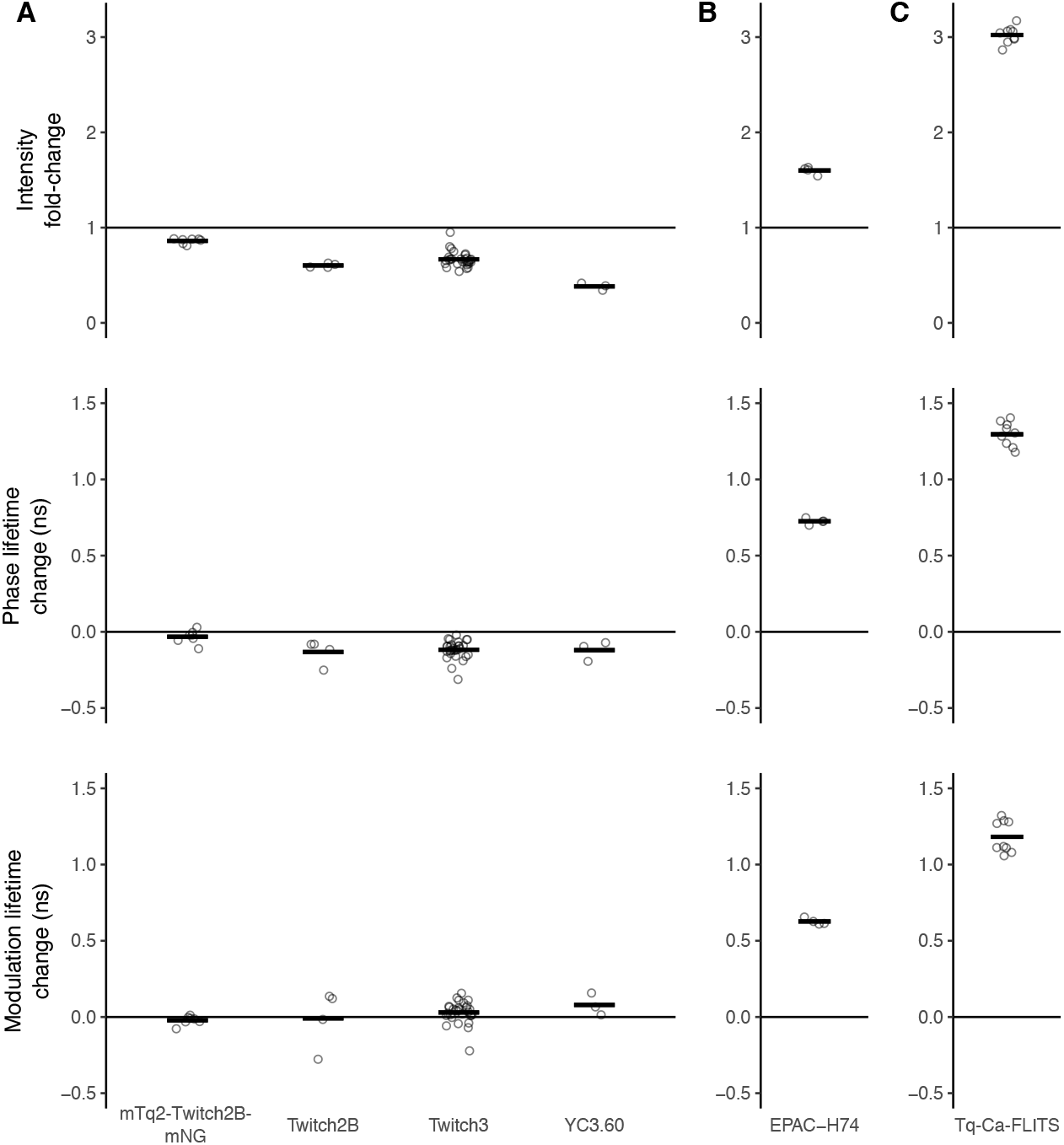
Intensity and lifetime contrast of biosensors. **A)** YCaM3.60, Twitch3, Twitch2B and mTq2-Twitch2B-mNG were transiently expressed in HeLa cells. Intensity and donor lifetimes were recorded before and after stimulation with 14 mM ionomycin combined with 5 mM CaCl_2_ (*n*=3 to 29). The intensity fold-change (top panel) and the absolute modulation and phase lifetime change (middle and bottom panel) of the donor fluorescent protein are plotted. **B)** Published donor intensity and lifetime change of EPAC-H74, a FRET-FLIM sensor for cyclic AMP^27^ (*n*=4) with a substantial lifetime contrast. **C)** For comparison we show the performance of new Turquoise calcium biosensor (Tq-Ca-FLITS). HeLa cells were transiently transfected with the sensor and stimulated with 14 mM ionomycin and 5 mM CaCl_2_ while imaging (*n*=9). Individual cell responses (circles) and their mean (line) are plotted for all sensors.

**Table S2.**
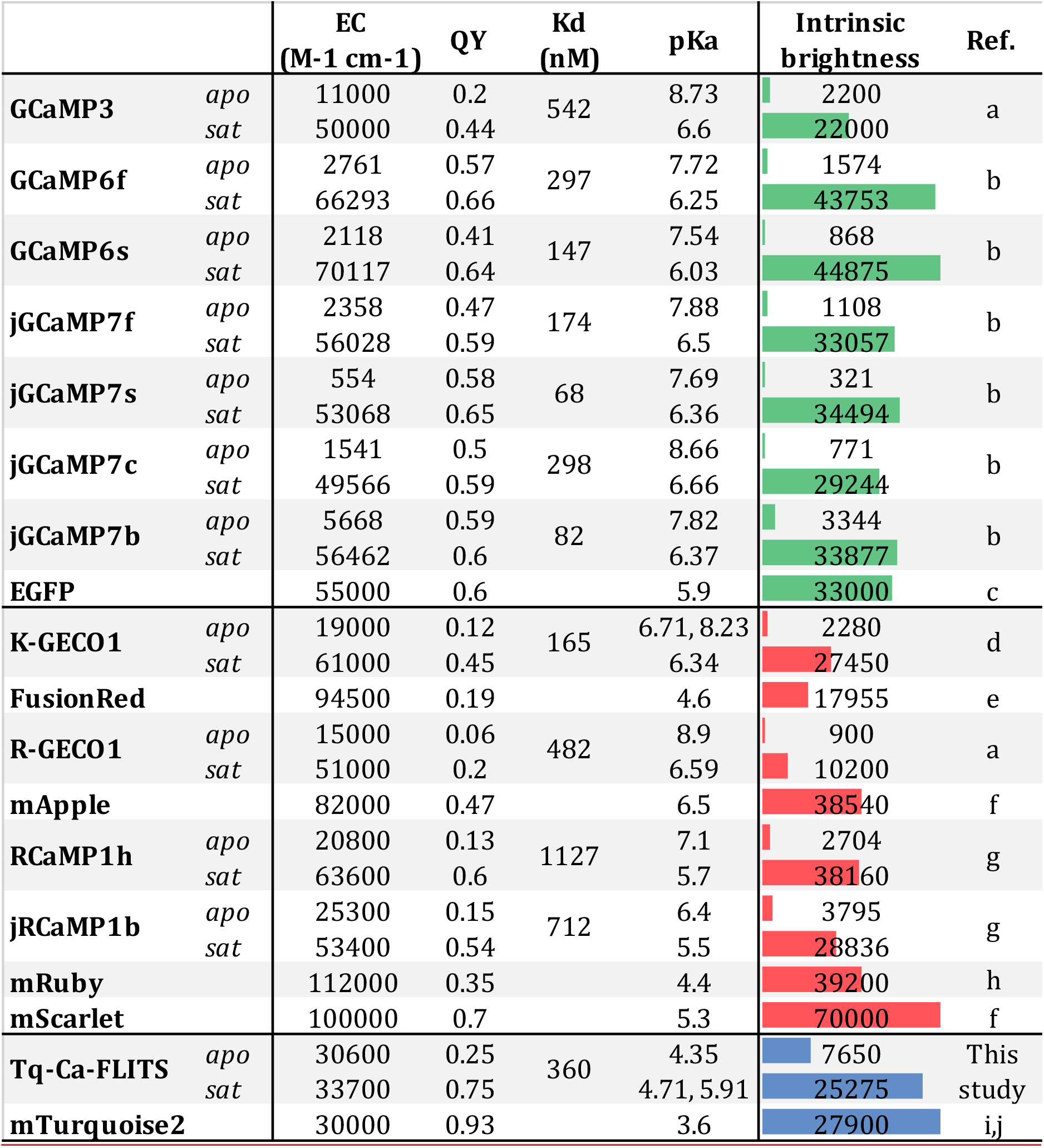
Properties of current intensity-based calcium sensors. Properties of the parent FP are also included: EGFP for all green sensors, FusionRed for K-GECO1, mApple for R-GECO1, mRuby for RCaMP1h and jRCaMP1b, mTurquoise2 for Tq-Ca-FLITS. mScarlet demonstrates the theoretical possibility for improvement of the brightness of the red sensors. Horizontal color bars indicate the relative intrinsic brightness compared to sensors and FPs of the same color (green, red or cyan). Tq-Ca-FLITS published here was added for comparison, which has a notably much higher relative intrinsic brightness in the calcium free state compared to other sensors. EC indicates the extinction coefficient, QY the quantum yield. References: a – Zhao et al. (2011)^6^, b – Dana et al. (2019)^3^, c – Patterson et al. (2001)^28^, d – Shen et al. (2018)^29^, e – Shemiakina et al. (2012)^30^, f – Bindels et al. (2017)^31^, g – Dana et al. (2016)^26^, h – Kredel et al. (2009)^32^, i – Goedhart et al. (2012)^33^, j – Cranfill et al. (2016)^34^.

|  |  | EC<br>(M<sup>-1</sup> cm<sup>-1</sup>) | QY | Kd<br>(nM) | pKa | Intrinsic<br>brightness | Ref. |
| --- | --- | --- | --- | --- | --- | --- | --- |
| **GCaMP3** | *apo* | 11000 | 0.2 | 542 | 8.73 | 2200 | a |
|  | *sat* | 50000 | 0.44 |  | 6.6 | 22000 |  |
| **GCaMP6f** | *apo* | 2761 | 0.57 | 297 | 7.72 | 1574 | b |
|  | *sat* | 66293 | 0.66 |  | 6.25 | 43753 |  |
| **GCaMP6s** | *apo* | 2118 | 0.41 | 147 | 7.54 | 868 | b |
|  | *sat* | 70117 | 0.64 |  | 6.03 | 44875 |  |
| **jGCaMP7f** | *apo* | 2358 | 0.47 | 174 | 7.88 | 1108 | b |
|  | *sat* | 56028 | 0.59 |  | 6.5 | 33057 |  |
| **jGCaMP7s** | *apo* | 554 | 0.58 | 68 | 7.69 | 321 | b |
|  | *sat* | 53068 | 0.65 |  | 6.36 | 34494 |  |
| **jGCaMP7c** | *apo* | 1541 | 0.5 | 298 | 8.66 | 771 | b |
|  | *sat* | 49566 | 0.59 |  | 6.66 | 29244 |  |
| **jGCaMP7b** | *apo* | 5668 | 0.59 | 82 | 7.82 | 3344 | b |
|  | *sat* | 56462 | 0.6 |  | 6.37 | 33877 |  |
| **EGFP** |  | 55000 | 0.6 |  | 5.9 | 33000 | c |
| **K-GECO1** | *apo* | 19000 | 0.12 | 165 | 6.71, 8.23 | 2280 | d |
|  | *sat* | 61000 | 0.45 |  | 6.34 | 27450 |  |
| **FusionRed** |  | 94500 | 0.19 |  | 4.6 | 17955 | e |
| **R-GECO1** | *apo* | 15000 | 0.06 | 482 | 8.9 | 900 | a |
|  | *sat* | 51000 | 0.2 |  | 6.59 | 10200 |  |
| **mApple** |  | 82000 | 0.47 |  | 6.5 | 38540 | f |
| **RCaMP1h** | *apo* | 20800 | 0.13 | 1127 | 7.1 | 2704 | g |
|  | *sat* | 63600 | 0.6 |  | 5.7 | 38160 |  |
| **jRCaMP1b** | *apo* | 25300 | 0.15 | 712 | 6.4 | 3795 | g |
|  | *sat* | 53400 | 0.54 |  | 5.5 | 28836 |  |
| **mRuby** |  | 112000 | 0.35 |  | 4.4 | 39200 | h |
| **mScarlet** |  | 100000 | 0.7 |  | 5.3 | 70000 | f |
| **Tq-Ca-FLITS** | *apo* | 30600 | 0.25 | 360 | 4.35 | 7650 | This study |
|  | *sat* | 33700 | 0.75 |  | 4.71, 5.91 | 25275 |  |
| **mTurquoise2** |  | 30000 | 0.93 |  | 3.6 | 27900 | i,j |

**Figure S3.**
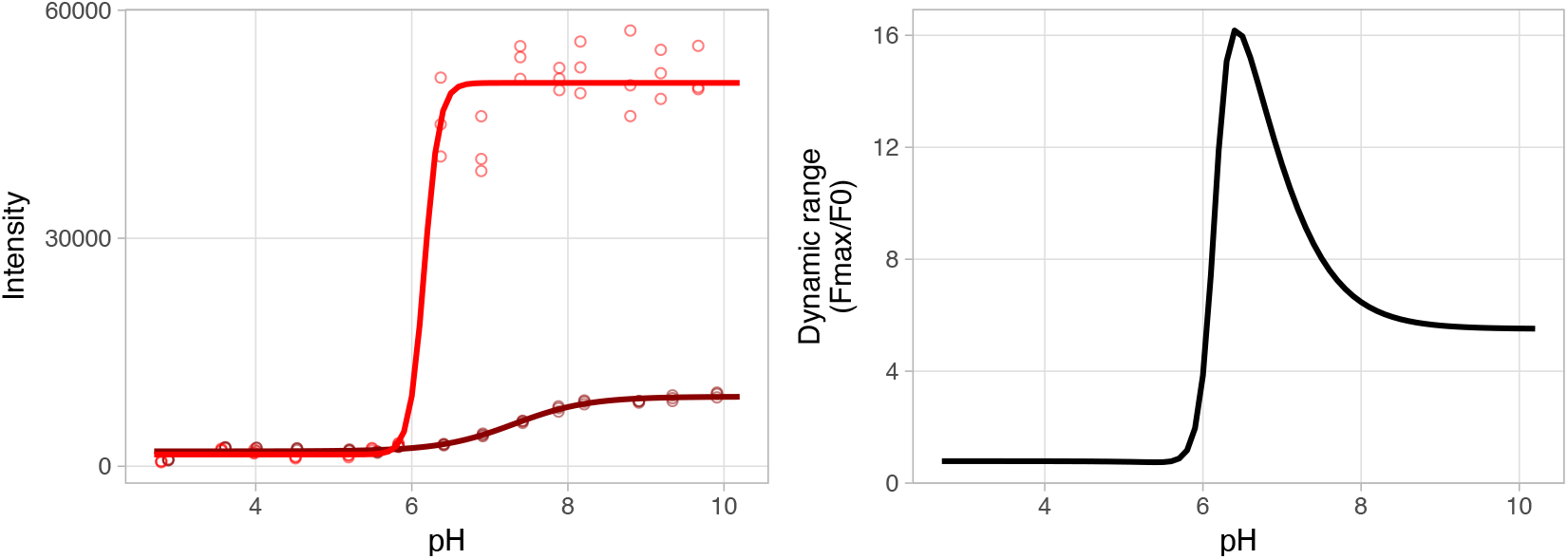
Intensity change of RCaMP1h as a response to pH. Left panel: A Hill curve with one p*K*_*a*_ value and Hill-coefficient was fitted through the measured data (circles, n=3) for the calcium bound (bright red) and unbound (gray) state, resulting in a pK_a,sat_ of 6.2 and a pK_a,apo_ of 7.3 Right panel: The calculated dynamic range was determined by division of the models fitted in the left panel.

Our aim was to engineer a new GECI using a single FP that displays a robust calcium-dependent lifetime contrast, with sufficient brightness to accurately measure calcium concentrations across its full dynamic range. We used mTurquoise2 as a template, since it has a high *QY*, high lifetime and we have a thorough understanding of residues that affects its *QY* and lifetime^33^. In addition, the p*K*_*a*_ of mTurquoise2 is low^34^ (∼3.6) and it is unlikely that a conformation change would affect the protonation state and *ε*.

Following the approach that led to the GECO series^6^, CaM and its binding peptide M13 were attached to new N- and C-termini of a circular permutated mTurquoise2 (cpTq2). Since information on circular permutated cyan fluorescent proteins is hardly available, we experimentally determined the optimal site for circular permutation of mTurquoise2. To this end, cpTq2 sensor variants were cloned into a new dual expression vector termed pFHL, which allows protein expression in mammalian cells and in the periplasm of *Escherichia coli*, and is suitable for isolation of the protein of interest (**Supplementary note 1, Figure S4**). Nine candidate sensors were constructed: seven with the new termini in the 7th β-strand and the remaining two in the 10th β-strand of the mTurquoise2 β-barrel (**Figure 1A**). Seven of the nine candidate sensors showed fluorescence in *E. coli* and HeLa cells. The sensor with the CaM and M13 attached to amino acids 149 and 150 respectively (which we named Tq-Ca-FLITS.0) showed about a 2-fold change in intensity in periplasmic fluid isolated from bacteria upon addition of calcium (periplasm test, **Figure S5A**), while the others showed little to no response. When expressed in HeLa cells, Tq-Ca-FLITS.0 showed a 3-fold intensity change upon addition of ionomycin (**Figure 1B and S5B**). We recorded the lifetime in both states of all seven candidate-sensors using frequency domain Fluorescence Lifetime Imaging Microscopy (FLIM). Tq-Ca-FLITS.0 again showed the largest response, with a striking phase and modulation lifetime change of over 1 ns (**Figure 1B-C and S5C**). The other variants showed only marginal changes in lifetime.

**Figure 1.**
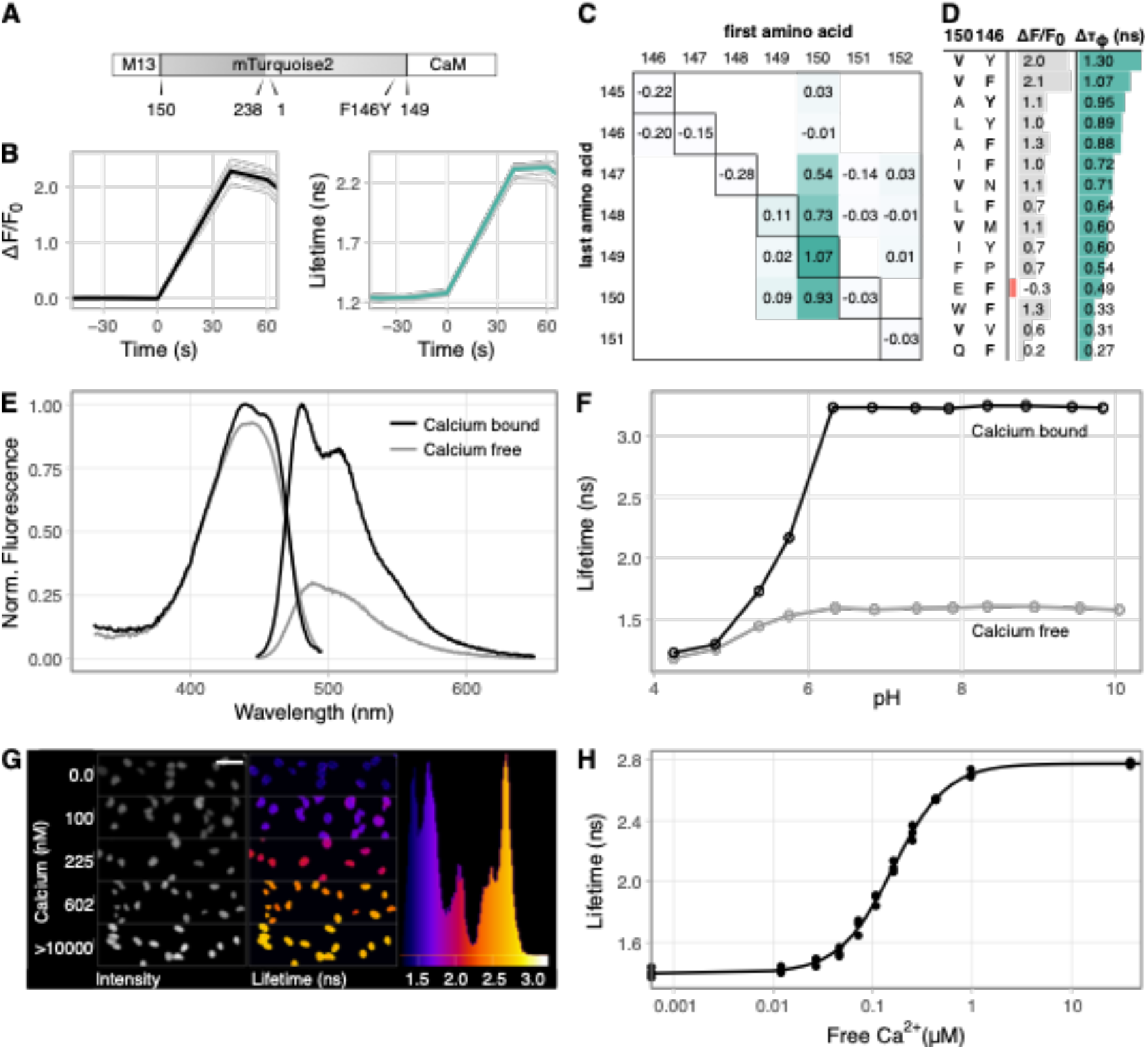
Development and characterization of the Turquoise calcium sensor. **A)** Schematic overview of the layout of Tq-Ca-FLITS, including key mutation F146Y. Positions of the original (amino acid 1 and 238) and the new (150 and 149) N- and C-termini are indicated. **B)** Intensity fold change over *F*_*0*_ and phase lifetime response of Tq-Ca-FLITS.0 in HeLa cells stimulated with ionomycin and calcium at *t*=0. Responses of individual cells (gray, *n*=7) and their mean (black or green) are plotted. **C)** Absolute phase lifetime change of various sensor variants in HeLa cells stimulated with ionomycin and calcium. The first amino acid of mTurquoise2 after the M13 peptide and the last amino acid before the CaM are indicated. On the diagonal axis (black outline) are the sensor variants that were created to find the ideal position to insert the CaM and the M13 peptide. The other variants contain 2 to 4 indels around the insertion site. Values are means of 2 to 23 individual cells. **D)** Intensity fold-change over *F*_*0*_ and absolute phase lifetime change of mutants of Tq-Ca-FLITS.0 in HeLa cells stimulated with ionomycin and calcium. Amino acids at position 150 and 146 of mTurquoise are indicated. The original residues (V and F) are printed in bold. Values are means of 2 to 10 individual cells. **E)** Normalized absorbance and emission spectra of Tq-Ca-FLITS *in vitro* for the calcium bound and unbound state. **F)** Tq-Ca-FLITS is pH insensitive from pH 6.2 onwards, as shown by the phase lifetime of the calcium bound and unbound states *in vitro* (*n*=3 with line average). **G)** The phase lifetime of Tq-Ca-FLITS stabilizes in HeLa cells during the *in vivo* calibration. The endpoint of 5 concentrations is shown. Visual display is generated by an ImageJ macro^_35_^. Scale bar is 50 μm. **H)** Calibration curve of the phase lifetime *in vivo*, with the calcium concentration on a logarithmic scale (*n*=3).

**Figure S4.**
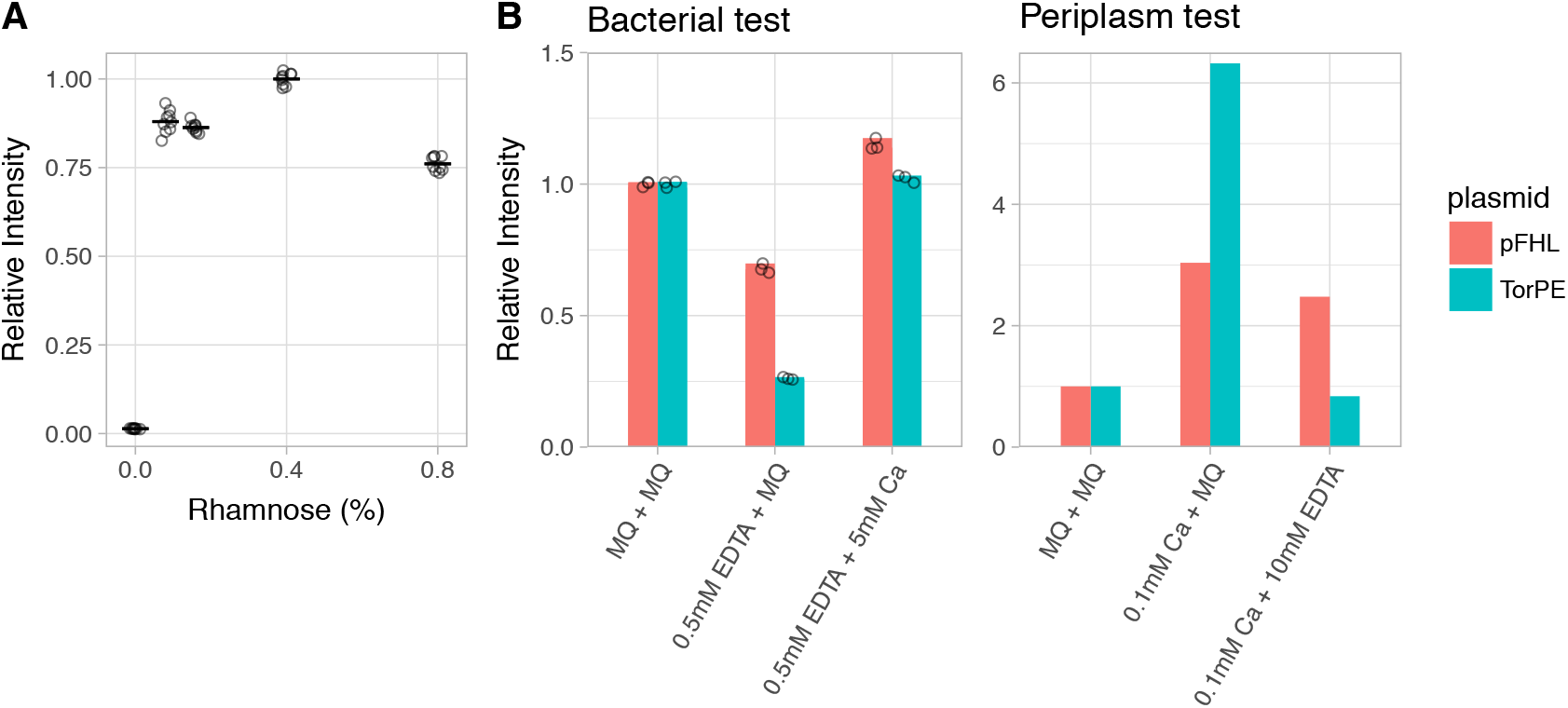
Verification of the new dual expression plasmid named pFHL. **A)** Normalized fluorescence intensities are shown of overnight grown *E. coli* cultures expressing R-GECO1 at different rhamnose concentrations, using the pFHL vector that contains a rhamnose promotor. Separate measurements (circles, *n*=9) and the mean are indicated. **B)** R-GECO1 responds to changing calcium concentrations in *E. coli* (Bacterial test, *n*=3) and in isolated periplasmic fluid (Periplasm test, *n*=1), by sequential addition of calcium and chelator EDTA. MilliQ water (MQ) was used a control. The performance of the pFHL vector in this test was compared to the pTorPE vector.

**Figure S5.**
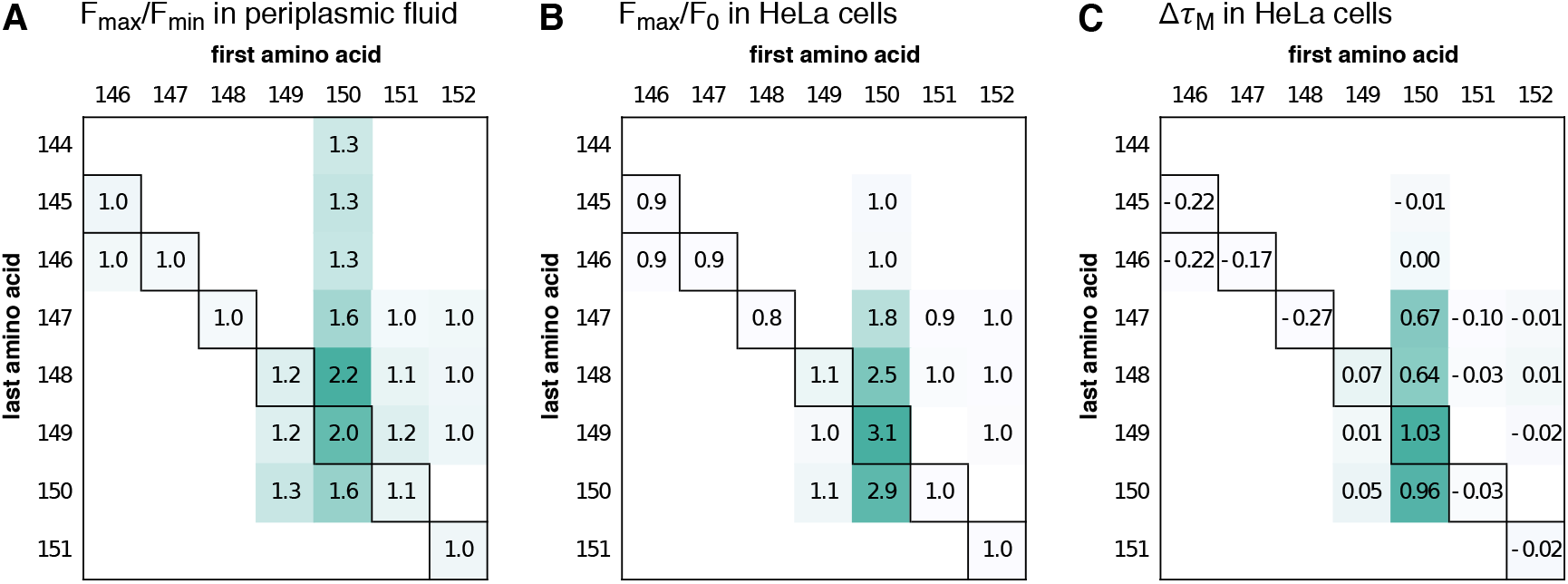
Screening the insertion site and the number of indels for the creation of Tq-Ca-FLITS. **A)** Intensity fold-change of different sensor variants in periplasmic shock fluid isolated from *E. coli*. **B)** Intensity fold-change and **C)** absolute modulation lifetime change (ns) in HeLa cells upon stimulation with ionomycin and calcium. In all panels are the first amino acid of mTurquoise2 after the M13 peptide and the last amino acid before the CaM indicated. See also the design of Tq-Ca-FLITS in **Figure 1A**. On the diagonal axis (black boxes) are the sensor variants that were created to find the ideal position to insert the CaM and the M13 peptide (mean of 8 to 23 cells). The other variants contain 2 to 5 indels around the insertion site (mean of 2 to 8 cells).

Next, we set out to improve our sensor by creating variants with up to 2 insertions or 5 deletions on both sides of the cpTq2, however this did not result in an improvement (**Figure S5**). Tq-Ca-FLITS.0 was therefore subjected to mutagenesis on two key residues that affect the fluorescence lifetime, i.e. F146 and V150 (original mTq2 numbering). F146 was previously shown to have a large influence on the lifetime of mTurquoise2^33^. V150 was selected for its position with respect to the chromophore, and in a small screen it showed to have an influence on the lifetime of mTurquoise2 (**Supplementary note 2, Figure S6**).

**Figure S6.**
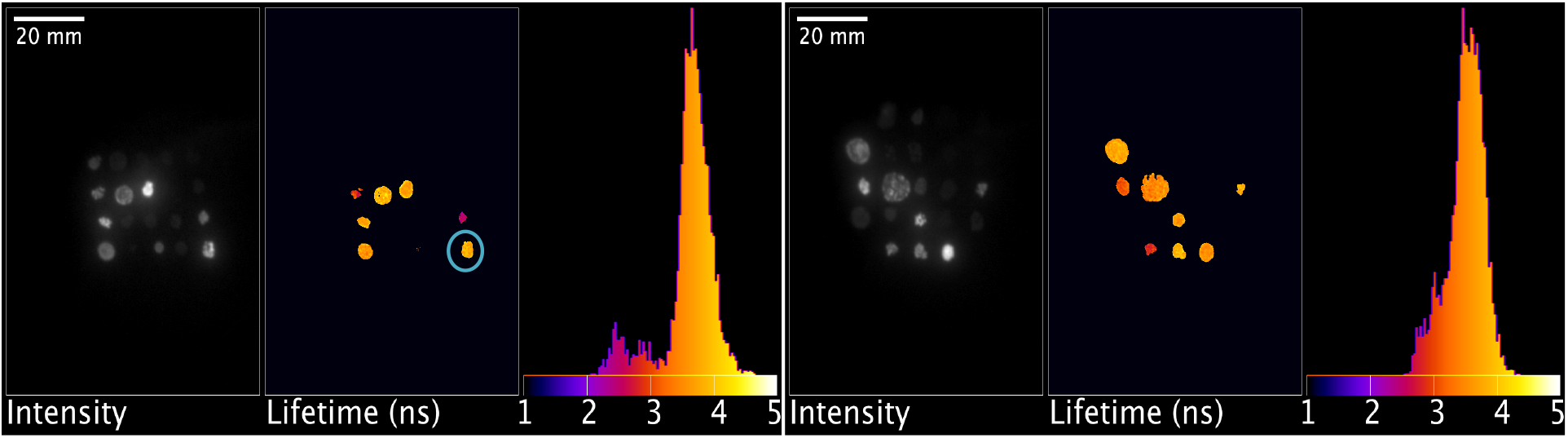
Lifetimes displayed by mTurquoise2 mutants in *E. coli* on agar plates. Intensity and modulation lifetime of mTurquoise2 mutated at position V150 are displayed. Two agar plates were imaged. A histogram of the modulation lifetimes is shown for each plate. A circle indicates non-mutated mTurquoise2. The panel was generated by an imageJ macro^35^.

A library of Tq-Ca-FLITS.0 containing F146X and/or V150X mutation(s) was initially screened in *E. coli* on agar plates, selecting for colonies showing a high intensity. We noticed that the sensors under these conditions are primarily in the high lifetime and high intensity state (**Supplementary note 3, Figure S7**). By selecting the high lifetime colonies we aimed to increase the overall brightness of the probe and increase the lifetime of the calcium bound state. Over 450 colonies were screened of which 60 ‘high’ intensity colonies were selected, and 17 ‘low’ and 16 ‘intermediate’ colonies as control. The selected colonies were screened in liquid bacterial culture, by monitoring the change in fluorescence intensity upon addition of calcium chelator EDTA to the culture. The periplasmic fluid of the best performing candidates was isolated for further testing (with the periplasm test) and their DNA sequence was determined.

**Figure S7.**
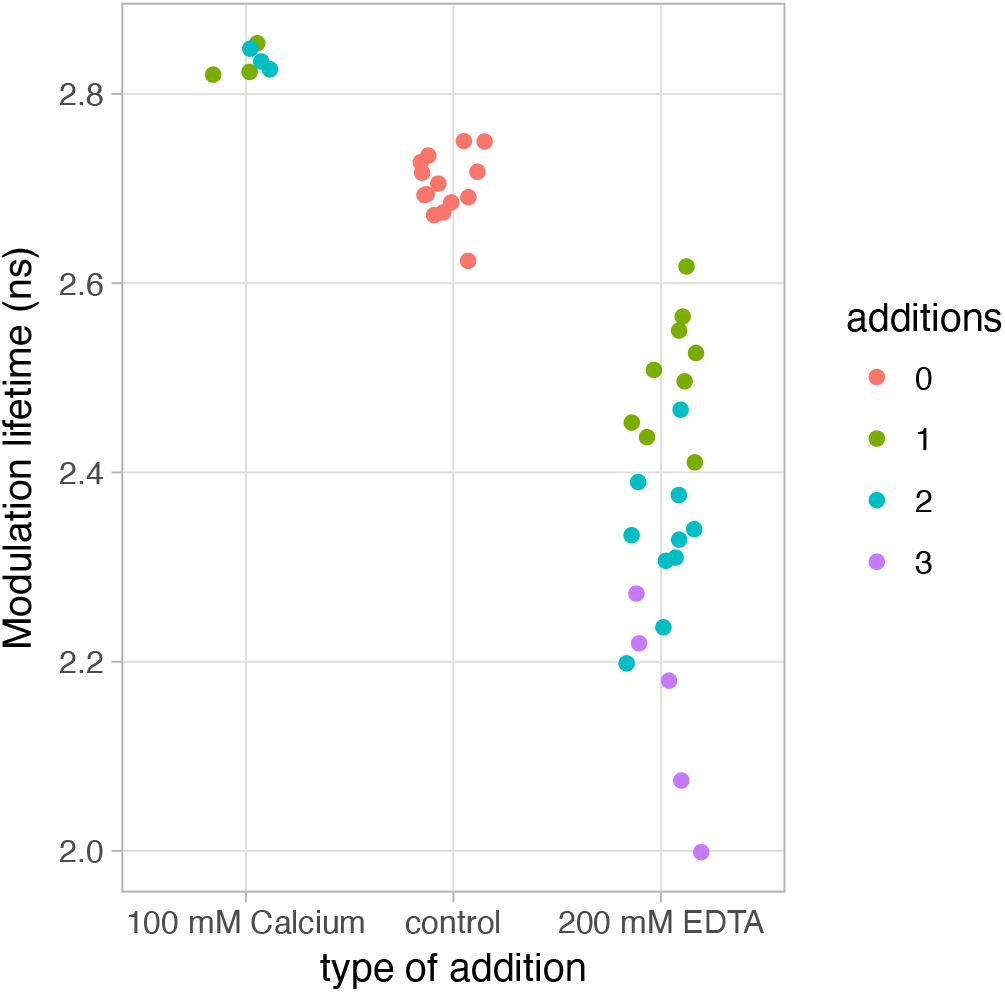
Modulation lifetime of bacterial colonies expressing Tq-Ca-FLITS.0. Colonies were stimulated 1, 2 or 3 times with a droplet of 100 mM calcium or 200 mM EDTA. Calcium increases the modulation lifetime and each drop of EDTA decreases the lifetime.

Optimal responses were observed when amino acids at position 150 were V or A, and F or Y at position 146 (**Figure S8)**. Based on these results five additional variants were constructed, with I, L or A at position 146 and F or Y at position 150. The designed variants and the top candidates from the screen were tested in HeLa cells for lifetime contrast (**Figure 1D**). We identified a new variant with a F146Y mutation that showed a comparable intensity response as the original variant and an increased phase lifetime response of about 1.3 ns. This variant was named Tq-Ca-FLITS, for Turquoise Calcium Fluorescence LIfeTime Sensor.

**Figure S8.**
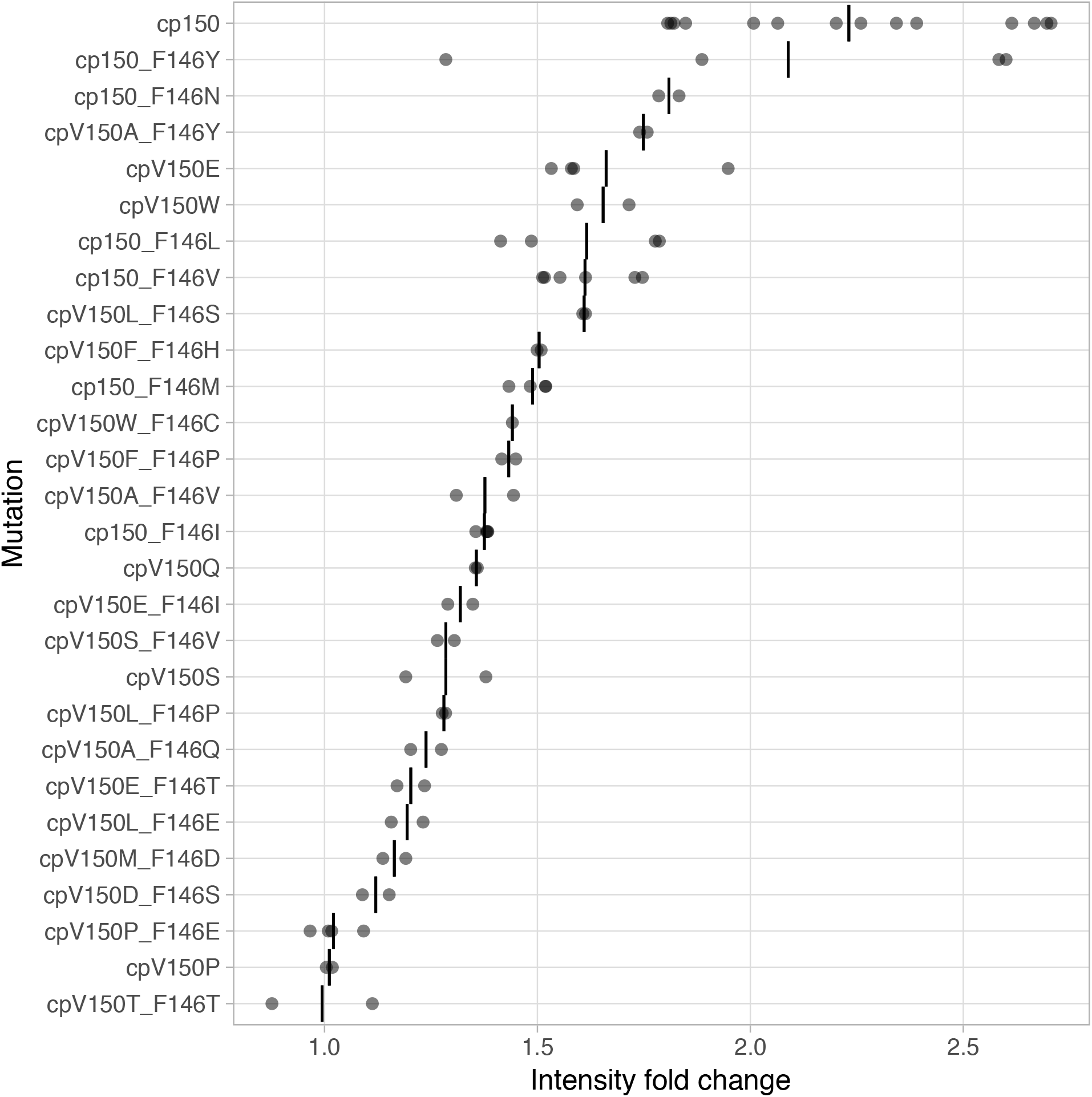
Intensity fold-change (*F*_*max*_/*F*_*0*_) of mutant sensors as measured by the periplasmic test. Gray circles indicate individual experiments and black lines indicate the means per mutant. The original amino acids at position 150 and 146 are V and F.

*In vitro* characterization of Tq-Ca-FLITS showed a substantial difference in quantum yield between the calcium bound and calcium free state (75% and 25% respectively), which is in line with the lifetime change. The extinction coefficient between the two states is comparable, unlike virtually all ‘GCaMPs’^3,23^ (**Table 1 and S3, Figure S9)**.

**Figure S9.**
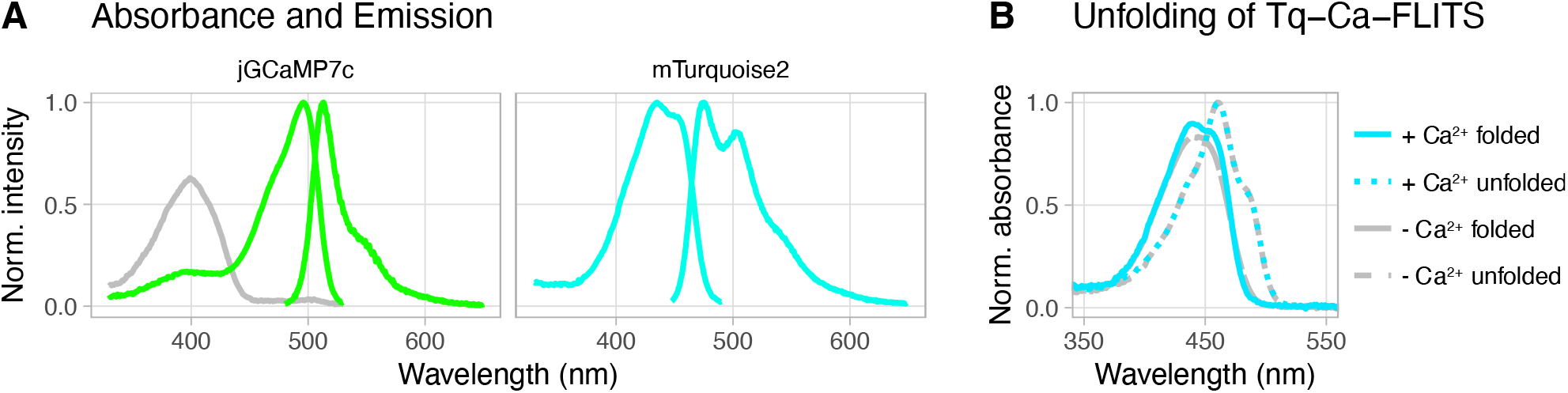
Properties of Tq-Ca-FLITS and jGCaMP7c *in vitro*. **A)** Absorption and excitation spectra of: jGCaMP7c (left panel) in the calcium bound (green) and unbound state (gray) and mTurquoise2 (right panel). **B)** Change in absorbance spectra of Tq-Ca-FLITS by unfolding the calcium bound (blue) and unbound (gray) state with 1 M NaOH.

**Table 1.** Properties of Tq-Ca-FLITS

|  | Spectral properties |  |  |  |  | Intensity data |  |  | Lifetime data |  |  |
| --- | --- | --- | --- | --- | --- | --- | --- | --- | --- | --- | --- |
| | $\lambda_{abs}$<br>(nm) | $\lambda_{em}$<br>(nm) | $QY$ | $\epsilon$<br>(mM<sup>-1</sup> cm<sup>-1</sup>) | $pK_a$ [n] | $K_d$ [n]<br>(nM)<br>pH 7.2 | $F_{max}/F_{min}$<br>*in vitro* pH 7.0 | $F_{max}/F_0$<br>*in vivo* | $\tau_\phi$<br>(ns) | $\tau_M$<br>(ns) | $K_d$ [n]<br>(nM) |
| Apo | 442 | 489 | 0.25 | 30.6 | 4.35 [0.86] | 360 [1.51] | 3.51 | 3.02 | 1.40 | 1.80 | 265 [1.63] |
| Sat | 439 | 481 | 0.75 | 33.7 | 1: 4.71 [0.70]<br>2: 5.91 [3.58] |  |  |  | 2.78 | 3.01 |  |

Spectral properties: $\lambda_{abs}$ – Absorbance maximum. $\lambda_{em}$ – Emission maximum. $QY$ – Quantum Yield relative to mTurquoise2. $\epsilon$ – Extinction coefficient at 440 nm. $pK_a$ [n] – Apparent $pK_a$ value, with the Hill coefficient [n] between brackets. A model with two $pK_a$ values was used for the calcium-saturated state.
Intensity: $K_d$ [n] – Apparent $K_d$ *in vitro*, with the Hill coefficient [n] between brackets. $F_{max}/F_{min}$ – ratio of maximum over minimum fluorescence intensity, or fluorescence of calcium bound state over calcium unbound state. $F_{max}/F_0$ – maximum intensity over starting intensity *in vivo*, as determined by stimulation with ionomycin and calcium.
Lifetime: quantitative *in vivo* readout with $\tau_\phi$ – phase lifetime. $\tau_M$ – Modulation lifetime. $K_d$ [n] – Apparent $K_d$ , with the Hill coefficient [n] between brackets, determined from the position of the lifetimes on a polar plot.

**Table S3.** Properties of Tq-Ca-FLITS compared to jGCaMP7c and mTurquoise2.

|  |  | Spectral properties |  |  |  |  |  | Sensor properties |  |  |
| --- | --- | --- | --- | --- | --- | --- | --- | --- | --- | --- |
| | | $\lambda_{abs}$<br>(nm) | $\lambda_{ex}$<br>(nm) | $\lambda_{em}$<br>(nm) | $QY$ | $\epsilon$<br>(M<sup>-1</sup> cm<sup>-1</sup>) | $pK_a$ [n] | $K_d$ [n] {CI}<br>(nM)<br>pH 7.2 | $F_{max}/F_{min}$<br>*in vitro* pH 7.0 | $F_{max}/F_0$ *in vivo* |
| Tq-Ca-FLITS | Apo | 442 | 446 | 489 | 0.25 | 30600 [440] | 4.35 [0.86] | 360 [1.51] {289-455} | 3.51 | 3.02 |
|  | Sat | 439 | 446 | 481 | 0.75 | 33700 [440] | 1: 4.71 [0.70]<br>2: 5.91 [3.58] |  |  |  |
| jGCaMP7c | Apo | 399 | n.d. | n.d. | 0.50<sup>\$</sup> | 1541** | 9.11 [0.60] | 337 [2.14] {309-369} | 127.8 | n.d. |
| | Sat | 496 | 500 | 513 | 0.59<sup>\$</sup> | 49566** | 6.82 [1.06] | | | |
| mTurquoise2 |  | 435 | 437 | 475 | 0.93* | 30000*<br>[434] | 3.67 [1.55] | n.a. | n.a. | n.a. |

Spectral properties: $\lambda_{abs}$ – Absorbance maximum. $\lambda_{ex}$ – Excitation maximum. $\lambda_{em}$ – Emission maximum. $QY$ – Quantum Yield relative to mTurquoise2. $\epsilon$ – Extinction coefficient, determined at the indicated wavelength. $pK_a$ [n] – Apparent $pK_a$ values, with Hill coefficient [n]. A model with two $pK_a$ values was used for the calcium saturated state of Tq-Ca-FLITS.
Sensor properties: $K_d$ [n] – Apparent $K_d$ , with the Hill coefficient [n] and the 95% confidence interval {CI} indicated. $F_{max}/F_{min}$ – ratio of maximum over minimum fluorescence intensity, or calcium bound state over calcium unbound state *in vitro*. $F_{max}/F_0$ – maximum intensity over starting intensity *in vivo*, as determined by stimulation with ionomycin and calcium ( $n=3$ ).
n.a./n.d.: not applicable/not determined.
\* Determined earlier in our lab<sup>33</sup>
\*\* Published by others<sup>3</sup>

**Figure S10.**
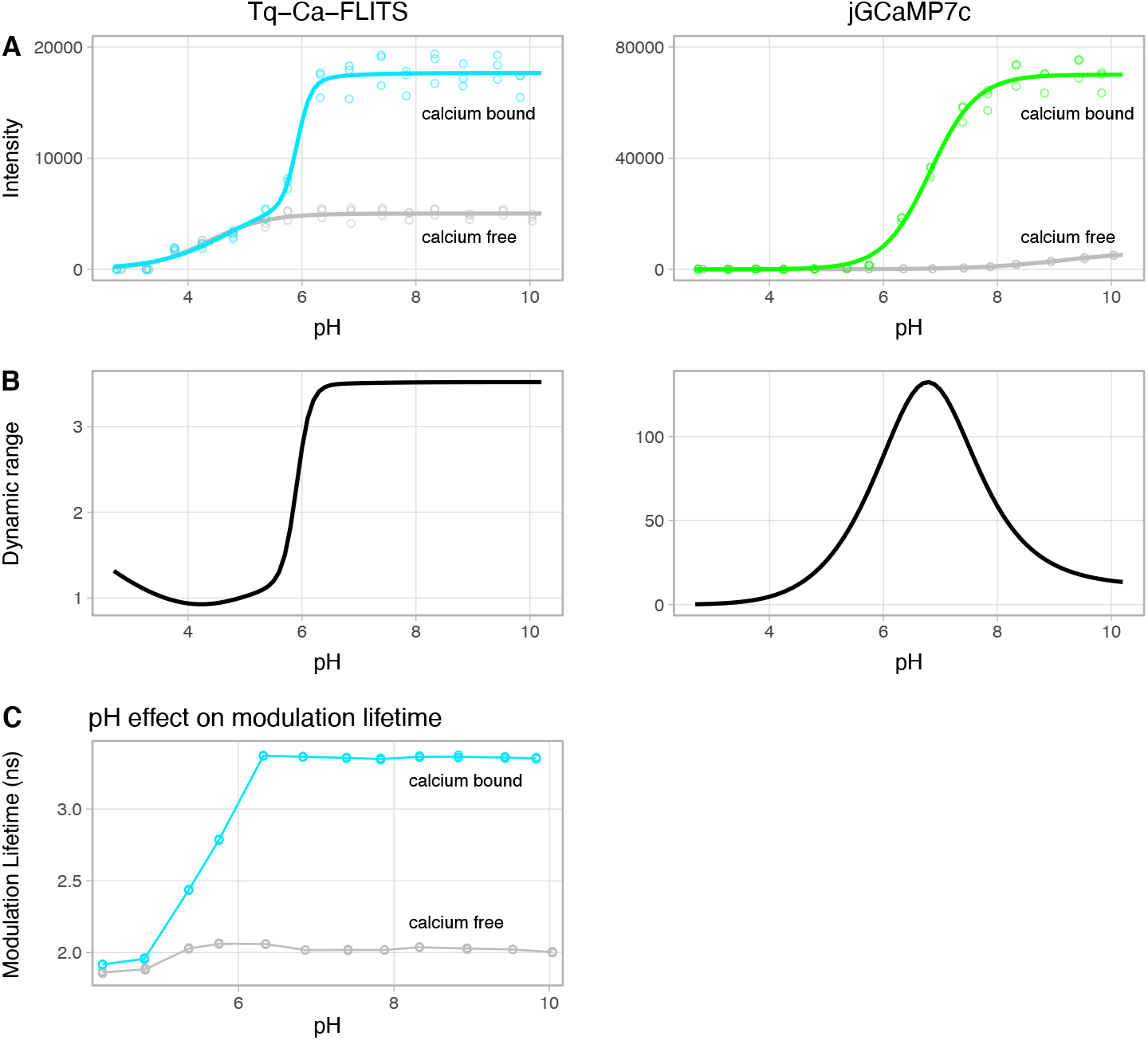
pH sensitivity of Tq-Ca-FLITS and jGCaMP7c *in vitro*. **A)** Intensity change of both sensors as a response to pH, for the calcium bound (blue/green) and unbound (gray) state (all *n*=3). A Hill curve (line) with one p*K*_*a*_ value and Hill-coefficient was fitted through the measured data (circles), except for calcium bound Tq-Ca-FLITS where a model with two p*K*_*a*_ values was used. **B)** The dynamic range of Tq-Ca-FLITS is stable in the biological pH range, while it fluctuates strongly for jGCaMP7c. The dynamic range is calculated as a ratio of the models presented in panel A. **C)** The modulation lifetime of Tq-Ca-FLITS is stable above pH 6.2 (*n*=3, with line average).

The pH and magnesium sensitivity of Tq-Ca-FLITS were investigated and compared to jGCaMP7c. Strikingly, both lifetime and intensity of Tq-Ca-FLITS are insensitive to pH above pH 6.2, making it a robust probe for biological measurements (**Figure 1F and S10**). Using the intensity data, we determined a p*K*_*a,apo*_ of 4.35 (Hill coefficient 0.86) for the free, unbound state of Tq-Ca-FLITS. A model with two p*K*_*a*_ values was used for the calcium bound state, and resulted in a p*K*_*a,sat,1*_ of 4.71 (Hill coefficient 0.70) and a p*K*_*a,sat,2*_ of 5.91 (Hill coefficient 3.58) (**Table 1 and S3**). The low and very similar p*K*_*a,apo*_ and p*K*_*a,sat,1*_ are likely a direct result of the pH dependency of the fluorescent protein in Tq-Ca-FLITS. The p*K*_*a,sat,2*_ probably shows the pH sensitivity of the four calcium binding domains of the CaM, which is also supported by the high Hill coefficient^36^. This p*K*_*a,sat,2*_ corresponds with the isoelectric point of CaM binding domains, which is shown to be around 6^37^. As expected, jGCaMP7c showed a clear pH sensitivity in the biological range.

We found a very low magnesium sensitivity for Tq-Ca-FLITS, both in intensity and lifetime readout (**Figure S11**). However, jGCaMP7c does show a marked change in dynamic range at the low magnesium concentrations (below 1 mM), which is reported to be right in the biological range^38^.

**Figure S11.**
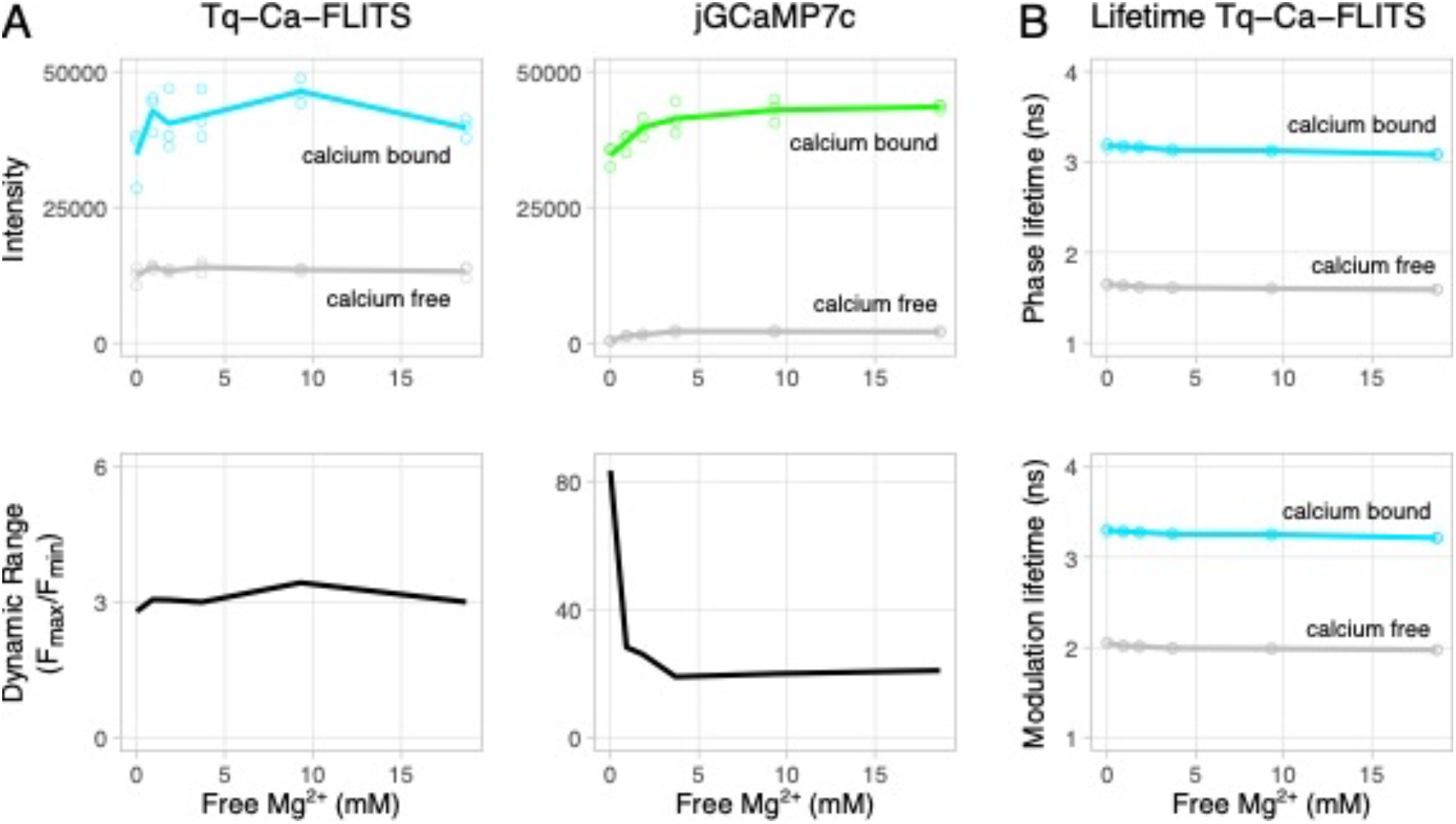
Magnesium sensitivity of Tq-Ca-FLITS and jGCaMP7c in vitro. **A)** Fluorescence intensity for the calcium bound (blue/green) and free (gray) state (top panels), and the corresponding dynamic range (bottom panels). **B)** Phase and modulation lifetime of Tq-Ca-FLITS in response to magnesium, for both the calcium bound (blue) and free (gray) state. Individual measurements (n=3) are indicated by dots, the line represents the average.

The intensity independent and quantitative readout of Tq-Ca-FLITS in combination with its specificity makes the sensor ideally suited for *in vivo* calibration^39^. To this end we established a HeLa cell line with stable expression of the sensor in the nucleus. Cells were incubated in calcium buffers and the calcium concentration in the cytoplasm was equilibrated with the outer environment by permeabilization of the membrane with digitonin. The concentration of calcium is plotted against the lifetime that was measured when equilibrium was reached (**Figure 1G-H, Movie S1**).

**Figure S12.**
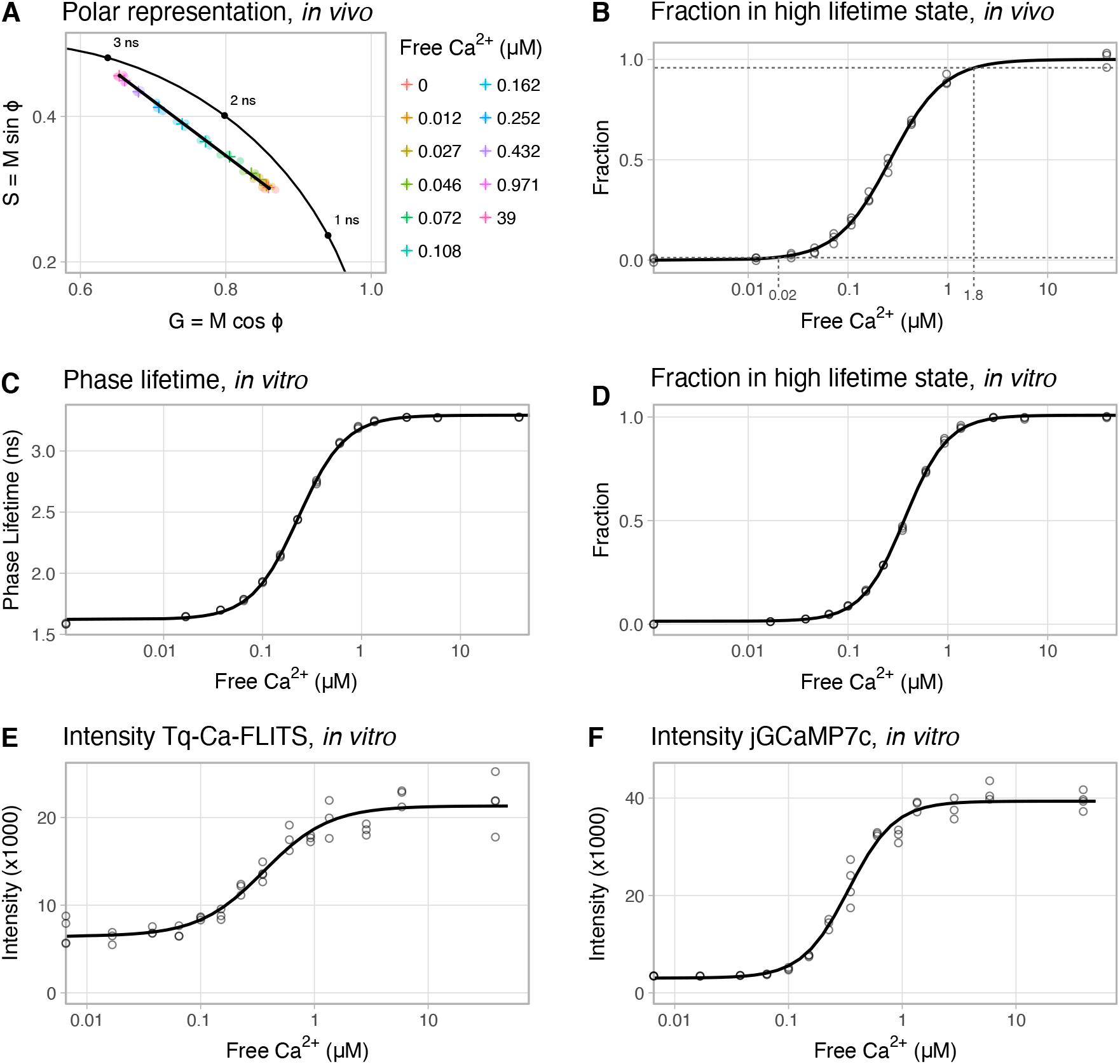
Calcium sensitivity of Tq-Ca-FLITS and jGCaMP7c. **A)** *In vivo* calibration of Tq-Ca-FLITS represented on a polar plot (*n*=3), where ‘*M*’ and ‘*Φ’* relate to the modulation and phase lifetime, respectively. Each dot represents the average of all pixels in a view with > 20 cells. The black line runs from the position of the calcium free state to the position of the calcium bound state. The fraction of sensor in the high state for each concentration was calculated from a projection on the black line, corrected for the intensity contribution of the two states. **B)** *In vivo* calibration of the fraction of Tq-Ca-FLITS in the high state as determined from a polar plot in panel A (n=3). The dotted horizontal lines indicate the borders of the measurable range, based on the 95% confidence interval of the fraction of the lowest and highest calcium concentrations. **C)** *In vitro* calibration of the phase lifetime of Ta-Ca-FLITS (*n*=3). **D)** *In vitro* calibration of the fraction of Tq-Ca-FLITS in the high state as determined from a polar plot (n=3). **E and F)** Sensitivity of the intensity readout of Tq-Ca-FLITS and jGCaMP7c *in vitro* (each *n*=3). In panels **B-F** the circles indicate individual measurements, and the line represents the fitted model. In all figures the calcium concentration is plotted on a logarithmic scale.

We converted the phase and modulation lifetime values of all calibration experiments to polar coordinates^40^, which resulted in a straight line on a polar (or phasor) plot (**Figure S12A**). For each calcium concentration we calculated the fraction of the sensor in the high lifetime state. This resulted in an apparent *K*_*d*_ of 265 nM, which is comparable to the GCaMP6 series^41^ (**Table 1 and S4, Figure S12B**). The apparent *K*_*d*_ *in vitro* was determined to be 372 nM, using isolated Tq-Ca-FLITS and applying a similar calculation on the lifetime data as for the *in vivo* calibration (**Table S4, Figure S12C-D**). When using the intensity data of the *in vitro* calibration, we found a similar apparent *K*_*d*_ of 360 nM (**Table 1 and S3, Figure S12E**). This is comparable to the apparent *K*_*d*_ *in vitro* of R-GECO1, the parent of Tq-Ca-FLITS^6^. Using the variation of the *in vivo* calibration we determined the detection range of the sensor to be 20 nM – 1.8 µM (**Figure S12B**).

**Table S4.** Lifetime properties of Tq-Ca-FLITS for quantitative measurements.

|  | *In vitro* calibration |  |  | *In vivo* calibration |  |  |
| --- | --- | --- | --- | --- | --- | --- |
| | $\tau_\phi$ {CI}<br>(ns) | $\tau_M$ {CI}<br>(ns) | $K_d$ [n] {CI}<br>(nM) | $\tau_\phi$ {CI}<br>(ns) | $\tau_M$ {CI}<br>(ns) | $K_d$ [n] {CI}<br>(nM) |
| Apo | 1.62<br>{1.61-1.64} | 1.97<br>{1.96-1.98} | 372 [2.03]<br>{364-380} | 1.40<br>{1.37-1.42} | 1.80<br>{1.75-1.86} | 265 [1.63]<br>{257-274} |
| Sat | 3.29<br>{3.28-3.30} | 3.35<br>{3.34-3.36} |  | 2.78<br>{2.74-2.81} | 3.01<br>{2.96-3.07} |  |

$\tau_\phi$ – phase lifetime. $\tau_M$ – Modulation lifetime. $K_d$ [n] – Apparent $K_d$ , with the Hill coefficient [n], determined from the position on a polar plot. 95% confidence intervals {CI} are indicated.

To enable detection in various cellular compartments, we generated probes that localize in the cytoplasm, at the Golgi apparatus or at the plasma membrane (**Figure S13**). Next, we examined the performance of the Tq-Ca-FLITS probe for quantitative intracellular calcium imaging in a number of biological systems. It has been well established that primary endothelial cells (ECs) respond to histamine with a transient intracellular calcium release^42^. We quantified the calcium levels after stimulation with 1 μM histamine and observed spatial heterogeneity of the calcium distribution, proving that Tq-Ca-FLITS is suitable for local quantification of intracellular calcium (**Figure 2A-B, Movie S2**).

**Figure S13.**
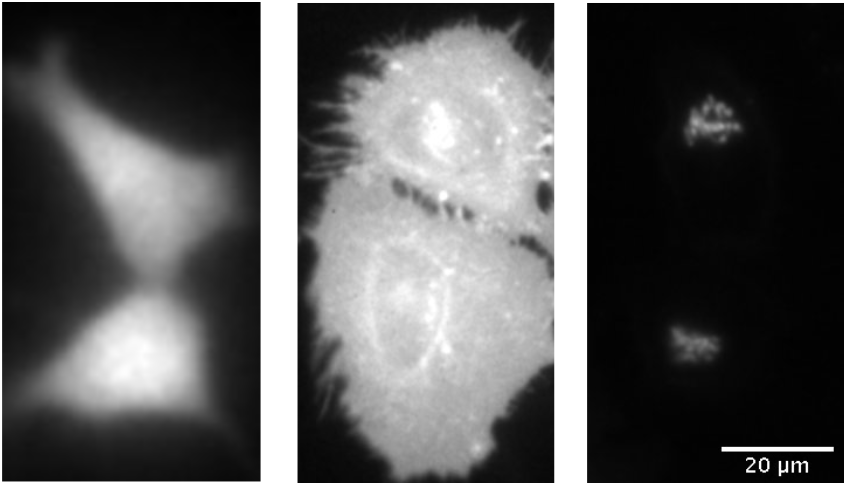
Organelle targeting of Tq-Ca-FLITS. The probe localizes to the cytoplasm without a targeting sequence (left panel), to the plasma membrane using an Lck-tag (middle panel), and the Golgi using a giantin-tag (right panel). All panels show two cells.

**Figure 2.**
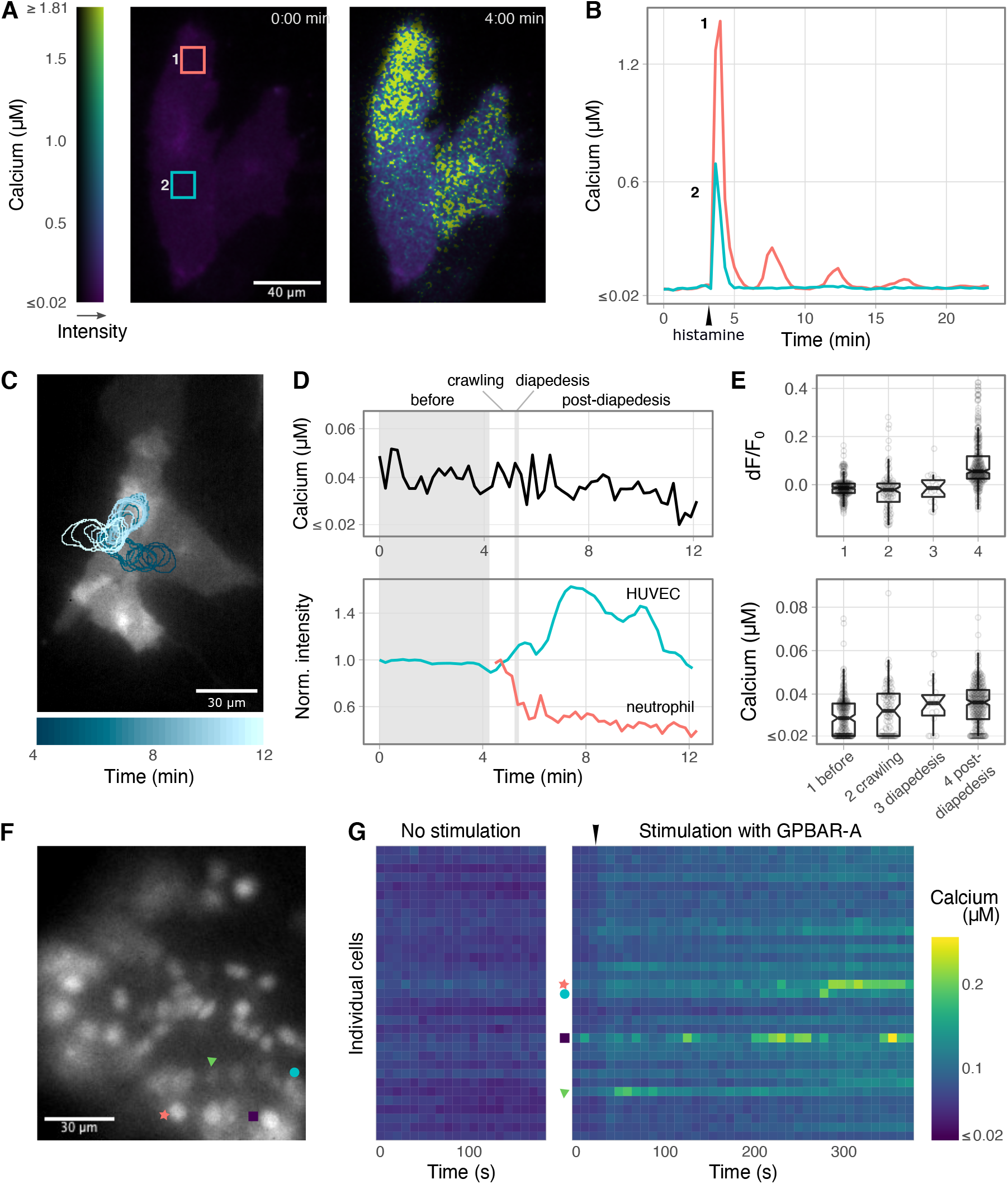
Quantitative calcium measurements with Tq-Ca-FLITS in primary endothelial cells and human organoids. **A)** Calcium levels in ECs with plasma membrane targeted Tq-Ca-FLITS before and after stimulation with 1 µM histamine, indicated by an arrow. The color scale indicates the concentration of calcium and the fluorescence intensity. Scale bar is 40 µm. **B)** Time trace of the regions indicated in panel A. **C-E)** Calcium concentration in ECs during TEM. The ECs express plasma membrane targeted Tq-Ca-FLITS and the leukocytes are labeled with Calcein Red-Orange. **C)** Fluorescence image of the ECs with the biosensor. The blue outline indicates the location of a neutrophil that crosses the EC monolayer. Scale bar is 30 µm. **D)** Top panel, calcium concentration in ECs measured at the location of the neutrophil indicated in panel C and bottom panel, the fluorescence intensity of Tq-Ca-FLITS and the leukocyte at the same location. The intensity of the neutrophil drops right after crossing the EC monolayer, while the intensity of Tq-Ca-FLITS increases. The concentration of calcium in the EC remains constant, as determined from the fluorescence lifetime of Tq-Ca-FLITS. **E)** Intensity fold change (dF/F_0_) of Tq-Ca-FLITS (top panel) and measured calcium concentration (bottom panel) in ECs before neutrophil contact, during crawling of the neutrophil, during diapedesis and post-diapedesis (16 TEM events, 7 measurements, 4 batches of neutrophils). The boxplot indicates the median, the 95% CI (notches), the first and third quartiles (hinges) and the 1.5x Interquartile Range (whiskers) All data points are also indicated by circles. **F-G)** Calcium changes measured in cells of a human small intestinal organoid stimulated with 10 μg/ml GPBAR-A. **F)** A human small intestinal organoid expressing Tq-Ca-FLITS in the nuclei of the cells. Scale bar is 30 µm. **G)** Heatmap of time traces of the calcium concentration of the cells depicted in panel F, without (left panel) and with (right panel) stimulation by GPBAR-A, indicated by an arrow. Markers indicate the corresponding cells in panel F.

We have previously shown that ECs actively prevent local leakage from blood vessels when leukocytes cross the endothelial barrier during transendothelial migration (TEM), by inducing a RhoA-dependent F-actin ring that serves as an elastic strap^43^. Whether calcium is involved in the subsequent pore closure is unknown. Although a role for calcium in TEM has been proposed, only the adhesion phase has been studied in some detail, with varying results^44^. The majority of studies used organic dyes with UV excitation and under non-physiological conditions, i.e. under the absence of flow (**Supplementary note 4**). We regarded conventional intensity-based GECIs unsuited for quantifying local calcium concentrations due to morphological changes of the cells, leading to substantial intensity changes unrelated to calcium concentrations.

We used a model system for TEM that utilizes flow to mimic physiological conditions, similar to the system we used previously to study RhoA activity^43^, but with improvements: (i) we used FLIM in combination with Tq-Ca-FLITS to measure a quantitative output and (ii) we simultaneously imaged fluorescence of the labeled leukocytes, enabling us to precisely correlate the different phases of TEM to the calcium concentration.

We achieved a temporal resolution of 13.5 seconds, which is sufficient to analyze calcium levels during TEM. Changes in the fluorescence intensity of both the endothelial cells and leukocytes (neutrophils) were observed (**Figure 2C-D, Movie S3**). The leukocyte intensity is affected by cell shape, as we are using a widefield microscope. High fluorescence intensity corresponds to a ball-like shape when the neutrophils are crawling, and low intensity is observed when the neutrophils spread out after completing diapedesis. The intensity of Tq-Ca-FLITS in the ECs was also affected by cell shape, and changes were observed when leukocytes continued to migrate under the EC monolayer. In contrast to the intensity changes, hardly any or no lifetime changes of the calcium probe were observed before adhesion, during crawling or during and after diapedesis of the leukocyte. This translates to hardly any or no calcium changes during TEM (**Figure 2E, Movie S3**). We analyzed a total of 16 tracks, capturing 98 crawling events and 19 diapedesis events. Also, the calcium concentration before adhesion (*n*=216) and post-diapedesis (*n*=278) was determined. The calcium levels in individual cells in almost all these events did not exceed 80 nM. This concentration is comparable to calcium concentrations reported in resting EC monolayers earlier^44–46^. In contrast, in the same events we did observe an intensity fold-change post-diapedesis (**Figure 2E**). The addition of ionomycin or histamine to the samples after the TEM assay showed a strong increase in intensity, lifetime and calcium concentration (> 1.5 µM), demonstrating that the calcium probe was fully functional in the experimental context (**Figure S14**). Our experimental approach that quantifies the calcium concentration in endothelial cells at different stages of TEM shows that calcium elevation (> 0.08 µM) in ECs is not essential for pore closure. Next to this, our data suggests that efficient crawling of neutrophils does not require an increase in baseline calcium levels.

**Figure S14.**
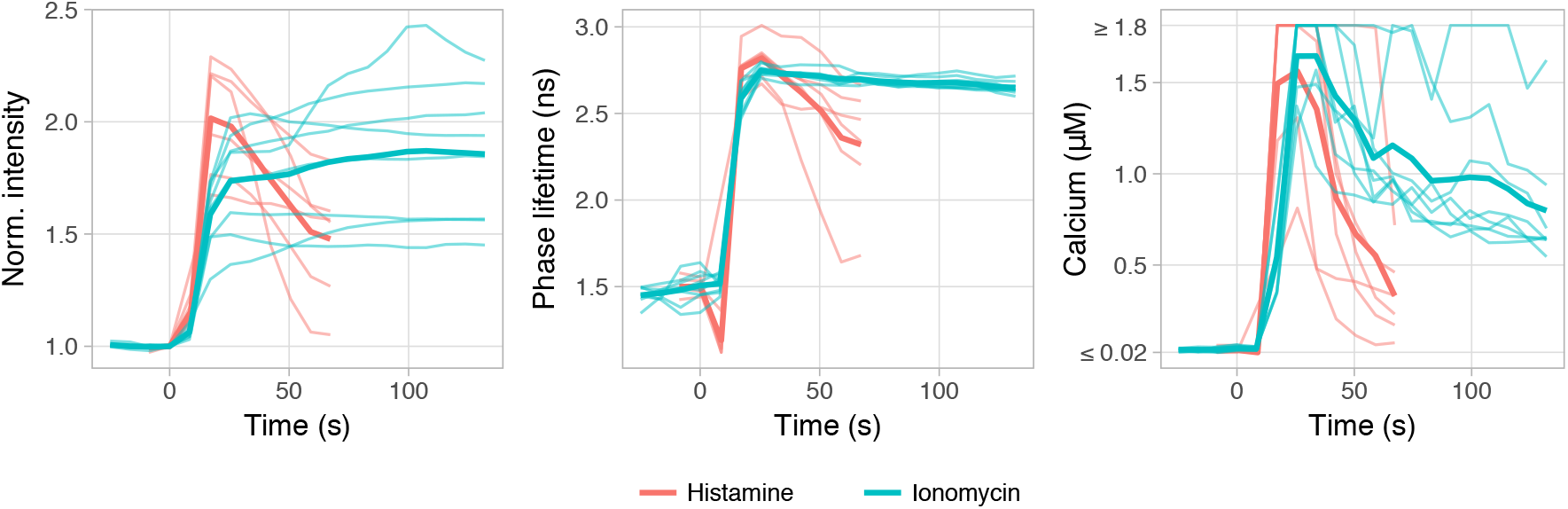
ECs expressing Tq-Ca-FLITS respond to histamine or ionomycin after the TEM assay. The change in intensity (left panel), change in lifetime (middle panel) and corresponding calcium concentration (right panel) are shown for individual cells with thin lines. Thick lines represent the mean. Tq-Ca-FLITS is fully functional in all measured cells (histamine *n*=6, ionomycin *n*=8).

Finally, the probe was used to quantify the calcium concentration in a human small intestinal organoid^47^. A nuclear-targeted Tq-Ca-FLITS variant was used to simplify the identification of individual cells. We previously observed an intensity change of the Tq-Ca-FLITS probe in response to stimulation of organoids with an odorant, however lifetime imaging was not performed in that study^47^. Here we investigated whether lifetime changes can be observed in this complex multicellular system. We differentiated organoids to the hormone-producing enteroendocrine cells (EECs), which express multiple G-Protein Coupled Receptors (GPCRs) that control secretion of their products^47^. *In vivo*, these cells represent less than 1% of the epithelium, but these can be enriched up to more than 50% in organoids. Calcium elevation is generally coupled to release of hormones from these cells. Lifetime imaging of the Tq-Ca-FLITS probe in nuclei of organoid cells revealed an intensity increase and a concomitant calcium increase in several cells when GPBAR-A was added. This drug is an agonist for GPBAR1, a GPCR that is expressed mainly by GLP-1-producing L-cells, a subtype of EECs, present in the organoid (**Figure 2F-G, Movie S4**). We observed a calcium increase by addition of GPBAR-A from 40-80 nM to an elevated baseline of 70-130 nM with occasional spikes up till 250 nM. As a control we saturated the calcium sensor in the organoid by addition of Triton-X100 and calcium to the medium (**Figure S15**).

**Figure S15.**
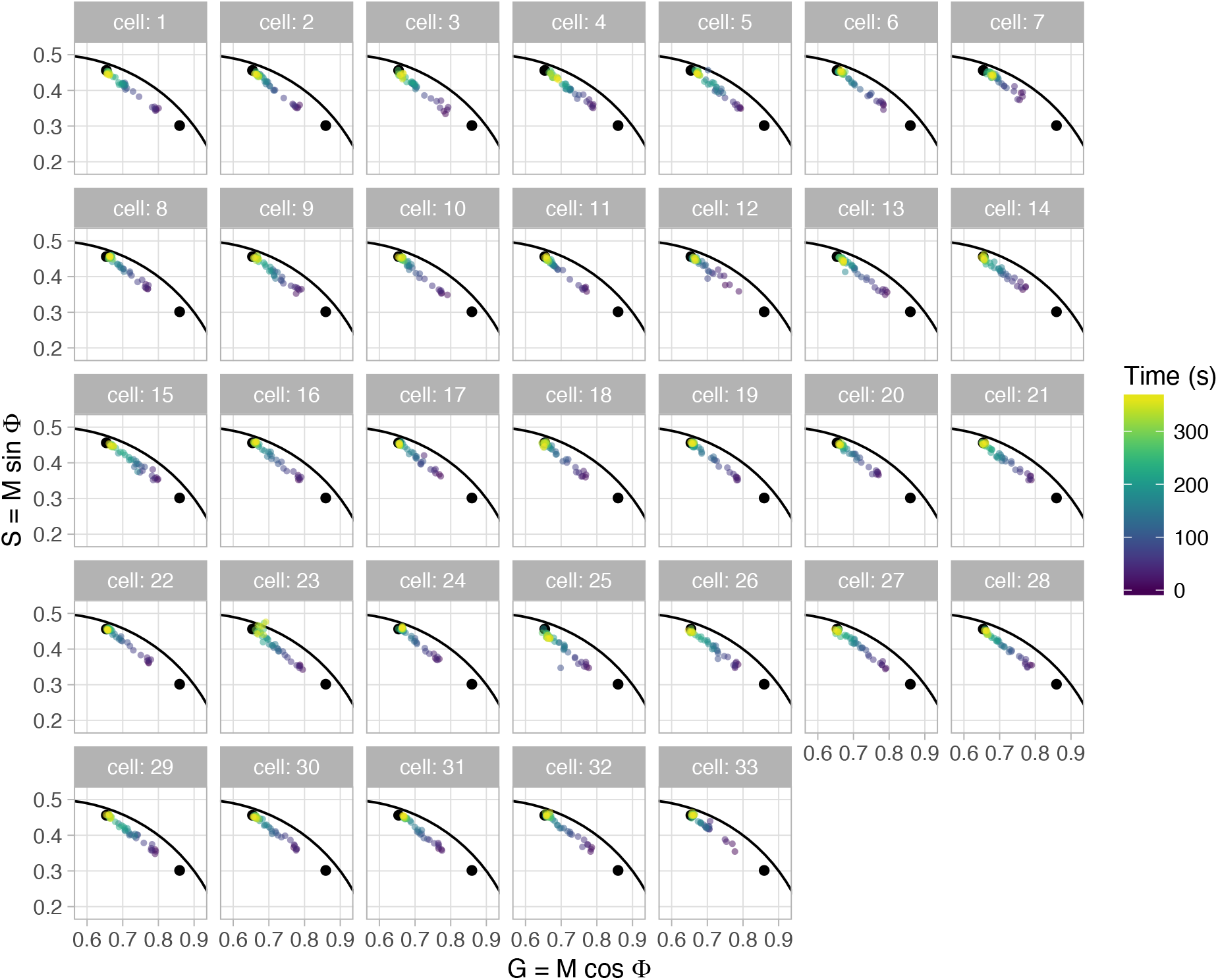
Tq-Ca-FLITS in human small intestinal organoid reacts to addition of 0.25 % (w/v) Triton-X100 and 12.5 mM calcium. A polar plot representation of the change in lifetime of individual cells (n=33) is given by the colored dots. The black dots indicate the positions of the calcium bound (upper left) and calcium free (lower right) states of the sensor. ‘M’ and ‘Φ’ relate to the modulation and phase lifetime, respectively. At the start of the measurement all cells show a lifetime near the calcium free state and towards the end of the measurement move completely (with a few exceptions) to the lifetime of the calcium bound state.

In summary, we have engineered a genetically encoded sensor for the quantitative detection of calcium. Our results demonstrate that a circular permutated variant of mTurquoise is a viable template for engineering sensors that (i) show lifetime contrast and (ii) are not sensitive to changes in the cytoplasmic pH range. The use of fluorescence lifetime to quantify calcium concentrations makes the measurements largely insensitive to changes in intensity. This simplifies the calibration in cells and results in a robust *in vivo* calibration. The lower calcium sensitivity of Tq-Ca-FLITS *in vivo* compared to *in vitro* is most likely caused by environmental differences. This demonstrates the need of an *in vivo* calibration when attempting quantification of intracellular concentrations. Unlike other GECIs, the Tq-Ca-FLITS is not optimized for intensity contrast and is therefore not the probe of choice for binary detection of calcium events. Instead, we have optimized this GECI for lifetime contrast, allowing simple, robust and precise quantification of intracellular calcium concentrations. We demonstrate accurate calcium imaging even when intensity fluctuations are caused by morphological changes, as observed during TEM. The temporal resolution was sufficient to capture the transmigration of neutrophil, however it could be greatly improved by choice of microscope. Our setup is not suitable for fast switching of the filters required for alternating imaging of cyan lifetime and red fluorescence, and it suffers from substantial response and dead times. When imaging only the lifetime in the CFP channel we reached a temporal resolution with Tq-Ca-FLITS of 3 seconds. Lifetime imaging can be further decreased to sub-second resolution by choice of improved and faster lifetime microscopy techniques such as a FALCON or Stellaris systems^48^ or siFLIM^49^. The relative intrinsic brightness of Tq-Ca-FLITS is 25-75% (‘off’- and ‘on’-state respectively) of mTurquoise2, the brightest cyan FP available. Moreover, in the ‘off’-state the intrinsic brightness of the Tq-Ca-FLITS is ∼76% of ECFP^50^, which is still commonly used in FRET ratio probes for timelapse imaging. Therefore, we expect that Tq-Ca-FLITS is suitable for faster and dynamic imaging of calcium concentrations than is demonstrated here.

The experiments with organoids show that we can accurately measure calcium levels in a complex 3D tissue. Human organoids are widely used for modeling of epithelial physiology. Previous work has highlighted – based on single cell transcriptomic analysis – the near-identical nature of organoid cells compared to their tissue counterparts^51^. The cellular heterogeneity of organoids may well facilitate quantifying cell type-specific calcium responses in a physiologically relevant context. We chose to assess calcium dynamics in human EECs, which eventually control hormone secretion. Human EECs differ greatly from their murine counterparts in terms of their expression profile of GPCRs, and therefore organoids represent a unique model to study EEC functioning in man. These endocrine cells represent important potential targets for treatments of metabolic diseases, as their hormones are involved in controlling key physiological processes such as appetite and insulin secretion. To increase the accuracy of measurements in thick samples, optical sectioning is necessary. This can be achieved by combing lifetime imaging with confocal scanning, light sheet imaging or multiphoton excitation.

To conclude, Tq-Ca-FLITS is the first GECI that incorporates a novel sensing mechanism based on a conformational change that directly modifies only the fluorescence quantum yield and fluorescence lifetime of a fluorescent protein independent of FRET (and without the need for a second fluorescent protein), providing contrast independent of sensor concentration. We anticipate that this novel sensor design can be easily combined with other sensor domains, e.g. reporting on phosphorylation and small molecule- or protein-binding, to generate an entire new class of fluorescence biosensors for the quantitative analysis of cellular processes.

## Methods

### General Cloning

We used *Escherichia coli* strain *E. cloni* 5-alpha (short: *E. cloni*, Lucigen corporation) for all cloning procedures. For DNA assembly, competent *E. cloni* was transformed using a heat shock protocol according to manufacturers’ instructions. For protein expression, *E. cloni* was grown using super optimal broth (SOB, 0.5% (w/v) yeast extract, 2% (w/v) tryptone, 10 mM NaCl, 20 mM MgSO4, 2.5 mM KCl) supplemented with 100 μg/ml kanamycin and 0.2% (w/v) rhamnose. 1.5% (w/v) agar was added for agar plates. Bacteria were grown overnight at 37 °C. Plasmid DNA was extracted from bacteria using the GeneJET Plasmid Miniprep Kit (Thermo Fisher Scientific) and the obtained concentration was determined by Nanodrop (Life Technologies).

DNA fragments were generated by Polymerase Chain Reaction (PCR), using Pfu DNA polymerase (Agilent Technologies) unless otherwise indicated. DNA fragments were visualized by gel electrophoresis on a 1% agarose gel, run for 30 min at 80 V. PCR fragments were purified using the GeneJET PCR purification Kit (Thermo Fisher Scientific) and digested with restriction enzymes to generate sticky ends. Restriction enzymes were heat inactivated at 80 °C for 20 min if necessary. Vector fragments were generated by restriction of plasmids and the correct bands were extracted from gel using the GeneJET Gel Extraction Kit (Thermo Fisher Scientific).

DNA fragments were ligated using T4 DNA ligase (Thermo Fisher Scientific), per the manufacturers’ protocol. Correct construction of plasmids was verified by control digestion and sequencing (primers 38-39, Macrogen Europe). All primers **(Table S5**) were ordered from Integrated DNA Technologies.

### Generating the dual expression vector

The pFHL-plasmid for dual expression was constructed using four DNA fragments: (i) the Kanamycin resistance gene and ColE1 origen of replication from a C1 plasmid (Addgene plasmid #54842), followed by (ii) the dual promoter region from a pDuEx plasmid (pDress were the mTurquoise2, large spatial linker and P2A sequences were removed from the plasmid using NheI restriction sites^35^), the sequence coding for R-GECO1 including TorA-tag from pTorPE-R-GECO1 (a gift from Robert Campbell (Addgene plasmid #32465; http://n2t.net/addgene:32465; RRID:Addgene_32465), and (iv) the terminator region from a pDuEx plasmid. The four DNA fragments were generated by PCR amplification with Phusion High-Fidelity DNA Polymerase (Thermo Fisher Scientific) (primers 1-8). The DNA fragments were assembled using Gibson assembly^52^. *E. cloni* was transformed with the Gibson mix and correct construction was verified by digestion analysis and sequencing of the plasmid.

### Engineering of Tq-Ca-FLITS

Circular permutated mTurquoise2 (cpTq2) variants were constructed by PCR amplification from a tandem construct containing two mTurquoise2 proteins connected by a flexible GGSGG-linker (primers 9-26). mApple in R-GECO1 on pFHL-R-GECO1 was replaced with different cpTq2 variants by digestion of the vector and PCR fragments with SacI and MluI, followed by ligation. To create a library of sensors with different linker lengths, a similar approach was taken, but now a library of cpTq2 fragments was generated by PCR using a mix of primers (primers 14-21 and 27). To create mutations at positions 146 and 150 of the fluorescent protein (mTq2 numbering), again the same approach was taken, now using primers containing altered or degenerated codons (primers 28-32).

Mutations were also made at position 150 of regular mTurquoise2, by PCR amplification with primers containing a degenerated codon (primers 33-34). The PCR mix was DpnI digested and used for transformation.

**Table S5.**
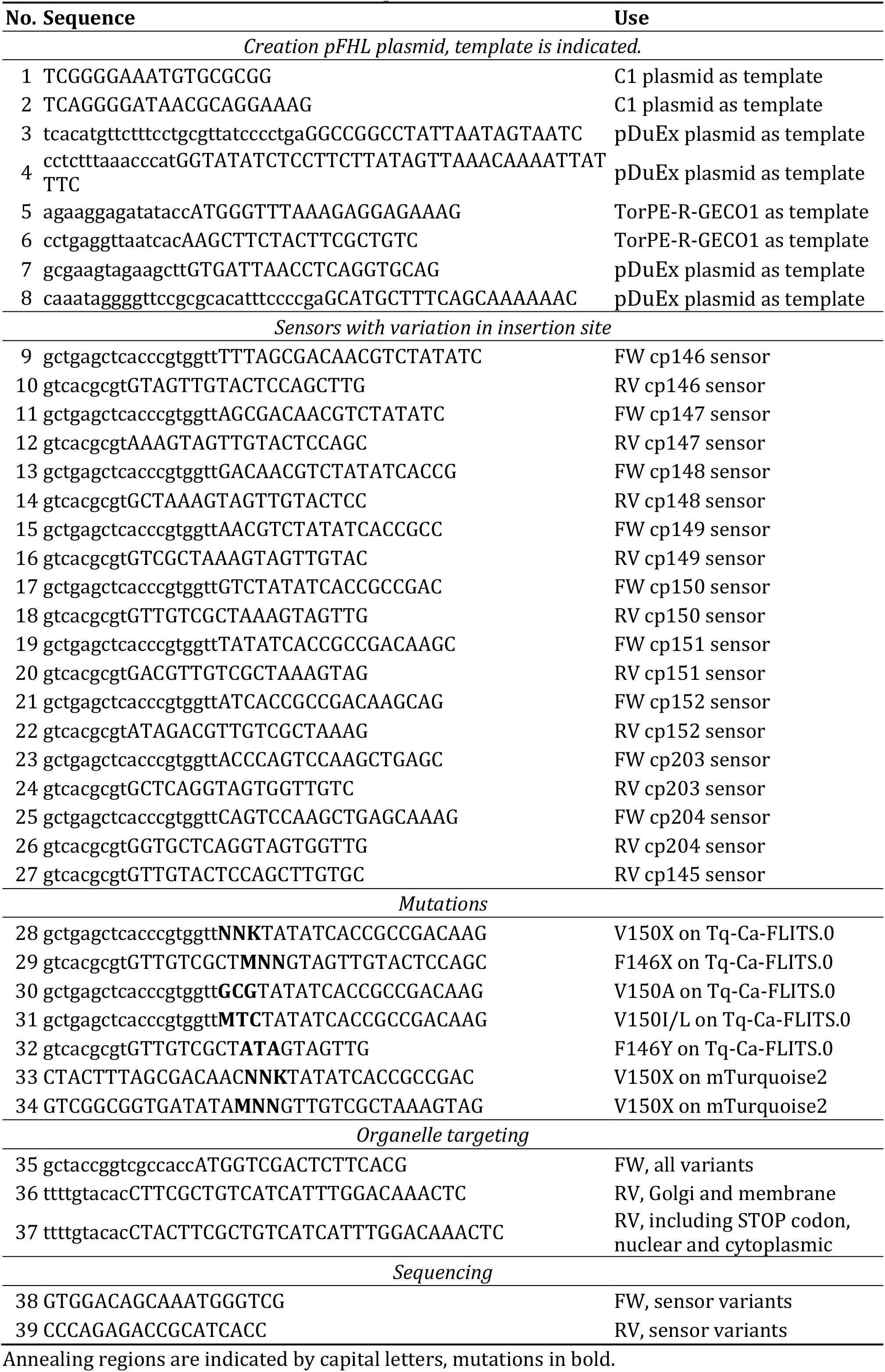
Primers for construction of Tq-Ca-FLITS.

### Plasmids for expression in mammalian cells

The sensor sequence was amplified by PCR (primers 35-37) to generate cytoplasmic, nuclear, membrane and Golgi targeted versions of Tq-Ca-FLITS for transient transfection of mammalian cells. DNA fragments and vectors carrying the desired tag were digested with AgeI and BsrGI, followed by ligation. Of the following vectors the fluorescent protein was exchanged for Tq-Ca-FLITS: 3xnls-mTurquoise2 (Addgene plasmid #98817) for a nuclear tag, pLck-mVenus-C1 (Addgene plasmid #84337) for a membrane tag, pmScarlet_Giantin_C1 (Addgene plasmid #85048) for a Golgi tag and mVenus-N1 (color variant of Addgene plasmid #54843) for an untagged version.

To generate stable cell lines, we used either the piggyBac transposase system or lentiviral transduction. To this end, Tq-Ca-FLITS including the 3xnls sequence was cloned into a PiggyBac vector containing a Puromycin resistance gene, by digestion with EcoRI/NotI and ligation, yielding pPB-3xnls-Tq-Ca-FLITS. The lentiviral plasmid for the doxycyclin-inducible expression of Tq-Ca-FLITS in organoids was adapted from the previously described pInducer20 x NLS-mKate2-P2A-HRAS^N17 53^. The following replacements were made via In-Fusion HD cloning kit (Clontech Laboratories) : NLS-mKate2 reporter fluorophore was replaced by PCR encoding NLS-mMaroon (primers FW 5’-3’ actagtccagACGCGtCcaccATGCcaaagaagaaacggaaggtaggatcaatggtgagcaagggcgag, RV 5’-3’ ttgtacagctccGTTAACccattaagtttgtgccccagtttgc); HRAS^N17^ was replaced by PCR encoding Tq-Ca-FLITS (primers FW 5’-3’ CCCTGGACCTGCTAGCatgggatcagatccaaaaaagaagagaaag, RV 5’-3’ gccctctagactcgagCTACTTCGCTGTCATCATTTGGACAAAC); ires-puro resistance cassette was replaced by ires-blast (primers FW 5’-3’ taaggatccgcggccGCATCGATGCCTAGTGCCATTTGTTCAGTG, RV 5’-3’ tctagagtcgcggccgcCATGCATTTAGCCCTCCCACAC). The resulting plasmid is indicated as pInducer-NLS-mMaroon-P2A-3xnls-Tq-Ca-FLITS.

We also constructed a plasmid for lentiviral transduction for normal expression of Tq-Ca-FLITS. Here In-Fusion Cloning was used to insert the coding sequences of H2B-mMaroon and 3xnls-Tq-Ca-FLITS connected with a P2A sequence in a lentiviral vector. The resulting plasmid is indicated as pLV-H2B-Maroon-P2A-3xnls-Tq-Ca-FLITS.

### Bacterial screening

*E. cloni* bacteria were used for two screening methods.

*(i) Bacterial test*. Bacteria expressing a sensor variant with the TorA-tag were grown overnight (O/N) in Luria-Bertani medium (LB, 10 g/L Bacto Tryptone, 5 g/L Bacto Yeast extract, 10 g/L NaCl). The bacterial suspension was pipetted in triplicate in a CELLSTAR 96-wells plate with black walls (655090, Greiner-Bio). Intensity was recorded at room temperature (RT) using a FL600 microplate fluorescence reader controlled by KC4(tm) software (Bio-Tek) with 430/25, 485/20 or 555/25 nm excitation and 485/40, 530/25 or 620/40 nm emission for respectively CFP, GFP or RFP, and averaging each well 10x. Intensity was again recorded after addition of 0.5 mM EDTA or MilliQ water (control). Intensities were divided by a well with 0.05 mg/ml Erythrosin B (EB), and a background (clear LB) was subtracted. Finally, wells were normalized to the first read of the same well to obtain the intensity fold-change *F*_*max*_/*F*_*0*_.

*(ii) Periplasm test*. Alternatively, the periplasmic shock fluid containing the sensor was isolated, using a cold osmotic shock protocol as described before^6^. Intensity of periplasmic fluid was measured before and after addition of 0.1 mM CaCl_2_ or MilliQ water, using the microplate reader described above. Intensities were divided by a well with 0.05 mg/ml EB, background (clear buffer) was subtracted and wells were normalized to the first read of the same well to obtain the intensity fold-change *F*_*max*_/*F*_*0*_. Each periplasmic isolate was measured in duplicate.

### HeLa cell culture

HeLa cells acquired from the American Tissue Culture Collection were maintained in full medium, DMEM + GlutaMAX (61965, Gibco) supplemented with 10% FBS (10270, Gibco), under 7% humidified CO_2_ atmosphere at 37 °C. Cells were washed with HBSS (14175, Gibco) and trypsinized (25300, Gibco) for passaging. No antibiotics were used unless otherwise stated. HeLa cells were grown on round cover slips (Menzel, no. 1, 24 mm diameter, Thermo Fisher Scientific) in a 6-wells plate for imaging. Transfection mixture was prepared in Opti-MEM (31985047, Thermo Fisher Scientific) with 2.25 μg Polyethylenimine in water (PEI, pH 7.3, 23966, Polysciences) and 250 ng plasmid DNA, and incubated for 20 min before addition to the cells. The coverslips were 1-or 2-days post-transfection mounted in an AttoFluor cell chamber (A7816, Thermo Fisher Scientific) and microscopy medium (137 mM NaCl, 5.4 mM KCl, 1.8 mM CaCl_2_, 0.8 mM MgSO_4_, 20 mM D-Glucose, 20 mM HEPES pH 7.4) was added.

### Stable expression of Tq-Ca-FLITS in HeLa

HeLa cells were transfected with pPB-3xnls-Tq-Ca-FLITS using PEI as transfection agent. One day post-transfection, transfected cells were selected by addition of puromycin (1 μg/ml) to the medium for 24h. The remaining cells were expanded for Fluorescence Assisted Cell Sorting (FACS). Briefly, cells were washed, trypsinized, resuspended in full medium and spun down at 1000 rpm for 4 min. Cells were washed twice in HF (2% FBS in HBSS) and resuspended in an appropriate volume of HF. The cell suspension was filtered through a 70 µm filter. Cells were sorted into full medium supplemented with P/S (100 U/ml penicillin and 100 μg/ml streptomycin) and 25 mM HEPES (pH 7.4) on a BD FACSARIA3, with 407 nm excitation and 502 nm long-pass and 510/50 nm band-pass emission filters. The sample chamber and collection devices were set at 4 °C for increased cell survival.

Single cells were gated based on Forward and Side Scatter (FSC/SSC). Live cell gating with DAPI was not possible due to spectral overlap with the fluorescence of Tq-Ca-FLITS. The positive gate for Tq-Ca-FLITS was determined based on untransfected wild-type HeLa cells. Cells were sorted into a high (34% of positive events) and low (66%) fluorescent pool. FACS data were analyzed with FlowJo. After sorting, cells were cultured with P/S for several weeks or until freezing down. Cells from the high pool were used for *in vivo* calibration of Tq-Ca-FLITS.

### Lifetime imaging

Fluorescence lifetime was recorded at RT with a 15-20 s interval, before and after addition of a mix of ionomycin (10 μg/ml, I-6800, LClaboratories) and calcium (5 mM). Two frequency domain FLIM microscopes were used.

(i) A home-build Zeiss setup controlled by Matlab 6.1 software, composed of an Axiovert 200M inverted fluorescence microscope (Zeiss) with a II18MD modulated image intensifier (Lambert Instruments) coupled to a CoolSNAP HQ CCD camera (Roper Scientific) and two computer-controlled HF-frequency synthesizers (SML 01, Rohde & Schwartz), one driving the intensifier and the other driving a 440 nm modulated laser diode (PicoQuant, LDH-M-C-440) through an MDL-300 driver unit^54^. The excitation light is modulated at 75.1 MHz and reflected by a 455 nm dichroic mirror onto the sample. Emission is filtered with a 480/40 nm band-pass emission filter. A 40X (Plan NeoFluar NA 1.3 oil) objective was used.

(ii) A LIFA setup composed of an Eclipse Ti microscope (Nikon) with a Lambert Instruments Multi-LED for excitation, a LI^2^CAM camera and a LIFA signal generator (all Lambert Instruments) to synchronize the light source and the camera. For CFP excitation a 446 nm LED was used, combined with a 448/20 nm excitation filter, a 442 nm dichroic mirror and a 482/25 nm band-pass filter. For RFP excitation a 532 nm LED was used, combined with a 534/20 nm excitation filter, a 561 nm dichroic mirror and a 609/54 nm band-pass filter (all filters from Semrock). Alexa488 or EB was used as a reference to calibrate the instrumentation, with a known mono-exponential lifetime of 4.05 ns^18,55,56^ or 0.086 ns^57–59^ respectively. Cells were imaged using a 40x (Plan Apo, NA 0.95 air) or a 60x (Plan Apo, NA 1.40 oil) objective.

Data from the Zeiss setup was analyzed as described before^60^. Data from the LIFA setup was converted to lifetime images by the LI-FLIM software (version 1.2.13). Regions of interest (ROIs) were selected to extract the average lifetime. Only cells with appropriate average intensity were selected to avoid influence of background fluorescence (> 200 for the Zeiss setup, > 2000 for the LIFA setup). The *in vivo* fold-change *F*_*max*_/*F*_*0*_ of Tq-Ca-FLITS was determined from the intensity data of a FLIM stack, corrected for background intensity.

### Ratiometric imaging

Fluorescent ratio imaging was with two different microscopes at 37 °C. (i) An Eclipse Ti microscope (Nikon) equipped with an Intensilight C-HGFIE (Nikon) for excitation and an Orca-Flash4.0 camera (Hamamatsu). Cells were imaged using a 40x (Plan Apo, NA 0.95 air) objective. For FRET imaging of YCaM3.60 we used a 448/20 nm excitation filter combined with a 4<u>42</u> nm dichroic mirror and a 482/25 nm emission filter for the donor, and a 514 nm dichroic mirror and a 542/27 nm emission filter for the acceptor. For MatryoshCaMP6s we used a 448/20 nm excitation filter, a <u>488</u> nm dichroic mirror and a 520/30 nm emission filter for imaging of the GFP and a 448/20 nm excitation filter, a 561 nm dichroic mirror and a 609/54 nm emission filter for LSSmOrange (all filters from Semrock).

(ii) An Axiovert 200M inverted fluorescence microscope (ZEISS) equipped with an Intensilight C-HGFIE (Nikon) for excitation and a CoolSNAP HQ CCD camera (Roper Scientific). Cells were imaged using a 40x (Plan Neofluar, NA 1.30 oil) objective. YCaM3.60 was excited with 420/30 nm. A 455 nm dichroic mirror was used. CFP fluorescence was collected at 470/30 nm and YFP fluorescence at 535/30 nm. MatryoshCaMP6s was excited with 440/30 nm followed by a 490 nm dichroic mirror. GFP fluorescence was collected at 525/40 nm and LSSmOrange fluorescence at 600/37 nm.

Background intensity was subtracted from the images and a ratio image was calculated using ImageJ (version 1.52k). The average intensity of each channel and the ratio was determined for individual cells.

### Protein isolation

His-tagged Tq-Ca-FLITS, jGCaMP7c, RCaMP1h and mTurquoise2 were isolated from bacterial culture essentially as described before, using Ni^2+^ loaded His-Bind resin^61^. In the final step, the isolated protein was overnight dialyzed in 10 mM Tris-HCl pH 8.0. No further purification was performed.

### Quantum yield

Purified Tq-Ca-FLITS and jGCaMP7c were diluted 10x in 10 mM Tris HCl with 100 µM CaCl_2_ or 5 mM EGTA, for the calcium bound or unbound state respectively. The absorbance spectra were measured with a spectrophotometer (Libra S70, Biochrom) between 260-650 nm for CFP and 260-700 nm for GFP (step size 1 nm, bandwidth 2 nm). Buffer without protein was used as reference. Each dilution was measured three times. Three dilutions were made using the initial dilution, with an absorbance at 440 nm (*A*_440_) of 0.002 < *A*_440_ < 0.02, each in triplicate. Emission and excitation spectra were recorded with a LS55 fluorimeter controlled by FL WinLab software (Perkin Elmer), with a step size of 0.5 nm and a scan speed of 200 nm/min and using buffer as reference. Emission was recorded at 450-650 nm (2.5 nm slit) with 440 nm excitation (4 nm slit). Excitation spectra were recorded at 250-490 or 250-530 nm (2.5 nm slit) measuring emission at 500 or 540 (4 nm slit), for CFP and GFP respectively.

Absorbance spectra were corrected by subtraction of the offset of the spectrum between 631-650 nm. Emission spectra were corrected for spectral sensitivity of the detector. The spectral area (*I*_*em*_) under corrected emission spectra was calculated by integration between 450-650 nm. The *A*_440_ was plotted versus the *‘I*_*em*_*’* and the slope *‘s’* was determined while forcing the regression line through the origin, *A*_440_ = *s* × *I*_*em*_. The Quantum Yield (*QY*) was determined using mTurquoise2 as reference with a know *QY* of 0.93 (**Equation 1**).

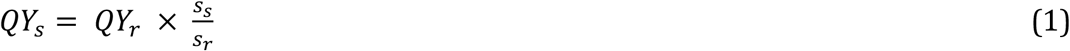

Subscripts ‘*s’* and ‘*r’* indicate the sample and the reference respectively. The average emission and excitation spectra were calculated from the highest protein concentration.

### Extinction coefficient

Purified Tq-Ca-FLITS was 4x diluted in calcium buffers containing 0 or 39 µM free calcium of the Calcium Calibration Buffer Kit #1 (C3008MP, Thermo Fisher Scientific). The absorbance spectra were measured before and > 5 min after addition of 1 M NaOH, at 260-650 nm with 1 nm step size and 1 nm bandwidth. Corresponding buffer was used as reference. The concentration of unfolded protein was determined using the Beer-Lambert law and assuming an extinction coefficient (*ε*) at 462 nm of 46 mM^-1^ cm^-1^ for the free cyan chromophore^62^. Next, *ε* was determined at 440 nm for the calcium free and bound states. The average absorbance spectra of three measurements were plotted.

### *In vitro* calibration

Purified Tq-Ca-FLITS and jGCaMP7c were diluted 100x in calcium buffers ranging from 0 to 39 µM, using the Calcium Calibration Buffer Kit #1 according to manufacturers’ instructions. Dilutions were made in triplicate. Fluorescence was measured at RT in a 96-wells-plate with black walls and flat glass bottom (89626, Ibidi) using a microplate fluorescence reader with settings as described under **Bacterial screening**. The fluorescence *‘f’* was fit to the Hill equation to determine the *K*_*d*_, using the Nonlinear Least Squares method of the R Stats Package (version 3.3.3) in R Studio (version 1.0.136) with default settings (**Equation 2**).

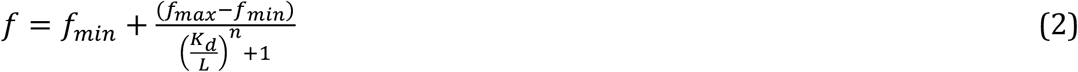

Subscripts *‘max’* and *‘min’* indicate the maximum and minimum fluorescence, *‘K*_*d*_*’* the microscopic dissociation constant, *‘L’* the known free Ca^2+^ concentration and *‘n’* the Hill-coefficient.

The lifetime of each well in the same 96-wells plate was recorded at RT using the LIFA setup described earlier, using the 40x (Plan Apo, NA 0.95 air) objective. Recorded sample stacks and a reference stack were converted into lifetime images by an ImageJ macro^35,63^.

The average phase and modulation lifetime (*τ*_*φ*_and *τ*_*M*_) of the full view were extracted. Both lifetimes were separately fitted with the Hill equation (**Equation 2**), with *‘τ’* instead of *‘f’*, using the Nonlinear Least Squares method of the R Stats Package (version 3.3.3) in R Studio (version 1.0.136) with default settings. The Phase (*Φ*) and modulation (*M*) were calculated from the recorded lifetimes and displayed in a polar plot as *G* and *S* coordinates (**Equations 3**).

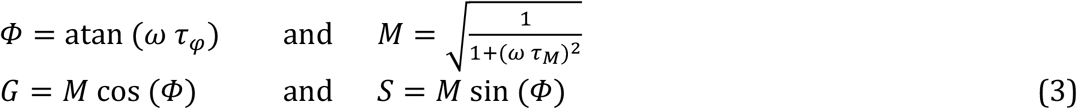

The angular frequency of modulation, *2πf*, is given by *‘ω’*. Measurements were projected on the straight line between the two extremes (*min* and *max*) and converted to line fraction *‘a’* (**Equations 4**).

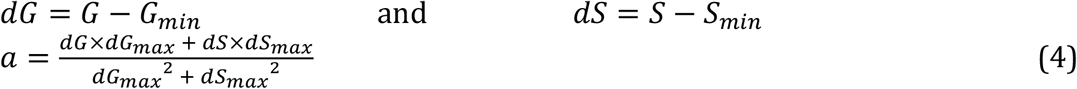

The line fraction was corrected of for the intensity contribution of the two states to find the true fraction *‘F’*, with *F*=1 representing all sensors in the calcium bound state (**Equation 5)**. The intensity ratio (*R*) between states *in vitro* was determined to be *R*=3.51 at pH 7.0, based on *f*_*max*_/*f*_*min*_ from the **pH sensitivity** experiments.

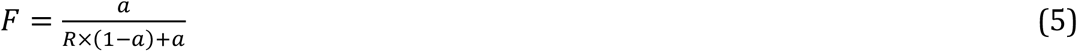

The fraction (*F*) was fitted with the Hill equation (**Equation 2**) with *‘F’* instead of *‘f’* to find the *in vitro K*_*d*_, using the Nonlinear Least Squares method of the R Stats Package (version 3.3.3) in R Studio (version 1.0.136), using the port algorithm.

### *In vivo* calibration

All steps were performed at 37 °C. Calcium buffers ranging from 0 to 39 µM (11 concentrations, in triplicate) were prepared using the Calcium Calibration Buffer Kit #1. HeLa cells stably expressing nuclear targeted Tq-Ca-FLITS were grown on cover slips and mounted in a cell chamber as described above. Cells were washed twice with HBSS without calcium (14175, Gibco) and calcium buffer was added. Cells were incubated for 15-20 min with 4 μg/ml rotenone and 1.8 mM 2-deoxy-D-glucose. Lifetime stacks were recorded using the LIFA setup described previously while adding 10 μM of digitonin. A 40x (Plan Apo, NA 0.95 air) objective was used and fluorescence lifetime was recorded every 20 s. Recorded data was converted into lifetime images by an ImageJ macro^35,63^.

The average lifetimes of all pixels in a view with intensity > 1000 were plotted over time (approximately 30-50 cells per view). Equilibrium was reached (reaction to digitonin) after > 6 min and the corresponding lifetimes were listed. Next, the same approach to determine the *in vivo K*_*d*_ was taken as for the *in vitro* data, using *R*=3.02 as determined from *F*_*max*_/*F*_*0*_ from HeLa cells stimulated with ionomycin and calcium.

Additionally, line fraction *‘a’* was fitted with the Hill equation (**Equation 2**), with *‘a’* instead of *‘f’*, to determine parameters for conversion of experimental data to calcium concentrations, without the need to correct for intensity contribution of the calcium free and bound states.

The sample standard deviation of the fraction *‘F’* was calculated for the highest and lowest calcium concentration, 0 and 39 µM. From this we determined the 95% confidence interval (*CI*). To determine the lowest fraction we can reliably measure, we added the 95% *CI* to the mean of the 0 µM calcium measurements. To determine the highest measurable fraction we did a subtraction.

### pH sensitivity

A series of buffers ranging from pH 2.8 to 10.0 were prepared, using 50 mM citrate buffer (pH 2.8-5.8), MOPS buffer (pH 6.3-7.9) and glycine/NaOH buffer (pH 8.3-10.0). Buffers additionally contain 0.1 M KCl and 0.1 mM CaCl_2_ or 5 mM EGTA. The pH of each buffer was determined including all components shortly before use. Purified Tq-Ca-FLITS, jGCaMP7c or RCaMP1h was diluted 100x in the prepared buffers.

Fluorescence was measured in triplicate in a 96-wells-plate with black walls and flat glass bottom (89626, ibidi) using a FL600 microplate fluorescence reader controlled by KC4(tm) software (Bio-Tek) with settings as described under **Bacterial screening**. The fluorescence intensity (f) was corrected for background and fit to the Henderson-Hasselbalch equation (**Equation 6**) using the Nonlinear Least Squares method of the R Stats Package (version 3.3.3) in R Studio (version 1.0.136), using the port algorithm restricted to *f min*≤0. The calcium bound state of Tq-Ca-FLITS did not fit to this model, therefore a model with two p*K*_*a*_ values was applied (**Equation 7**).

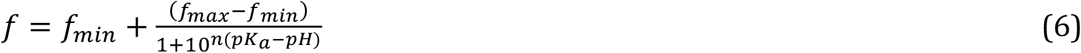

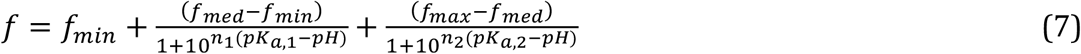

The *‘n’* indicates the Hill coefficient, *‘*p*K*_*a*_*’* the apparent p*K*_*a*_ and *‘min’, ‘med’* and *‘max’* the minimum, a medium and the maximum fluorescence. The model for the Ca^2+^ saturated state was divided over the model of the Ca^2+^ free state to gain the dynamic range.

The lifetime of each well in the same 96-wells plate was recorded at RT similar to the *in vitro* calibration. Recorded data was converted into lifetime images by the LI-FLIM software (version 1.2.13). The average phase and modulation lifetime (*τ*_*φ*_ and *τ*_*M*_) of the full view were extracted.

### Magnesium sensitivity

A series of buffers was prepared at pH 7.1 and 20 °C, each designed to contain a specific concentration of free magnesium ions (0, 0.9, 1.8, 3.7, 9.3 and 18.7 mM) and either 0 or 1 mM free calcium, as calculated using the program ‘*Ca-Mg-ATP-EGTA Calculator v1*.*0 using constants from NIST database #46 v8’*. The buffers including calcium contain each 1 mM CaCl_2_, 20 mM HEPES pH7.1 and 0, 0.9, 1.8, 3.7, 9.3 or 18.7 mM MgCl_2_. The calcium free buffers contain each 3 mM EGTA, 20 mM HEPES pH7.1 and 0, 1, 2, 4, 10 or 20 mM MgCl_2_. The expected trace of calcium in the calcium free buffers is 1 nM or less. The pH was measured after mixing of the buffers, and checked again before use.

Purified Tq-Ca-FLITS and jGCaMP7c were 100x diluted in the buffers and the intensity was recorded similar as done for the pH measurements. The dynamic range was calculated as the average intensity of the calcium bound state divided by the calcium free state. The lifetime of Tq-Ca-FLITS was recorded and processed in the same manner as described under **pH sensitivity**.

### Endothelial Cell (EC) culture and transendothelial migration (TEM)

Primary Human Umbilical Vein Endothelial Cells (HUVECs) acquired from Lonza (P1052, Cat #C2519A) were maintained in culture flasks pre-coated with fibronectin (FN, 30 μg/mL, Sanquin) in EGM-2 medium supplemented with SingleQuots (CC-3162, Lonza) and P/S, under 5% humidified CO_2_ atmosphere at 37 °C. HUVECs were passaged by washing twice with phosphate buffered saline (PBS) and trypsinization. Trypsin was inactivated with Trypsin Neutralisation Solution (CC-5002, Lonza). HUVECs were transfected by microporation at passage #4 with 2 μg plasmid DNA containing the Lck version of Tq-Ca-FLITS. For microporation the Neon Transfection System (MPK5000 Invitrogen) and corresponding Neon transfection kit were used according to manufacturers’ protocol. We used the R buffer from the kit, the 100 μL tips and a 30 ms pulse of 1300 V. After microporation, HUVECs were directly seeded on FN-coated round cover slips for imaging, similar as described for HeLa cells. For TEM experiments, the buffer was removed by centrifugation (200 g, 3 min, RT) and HUVECs were seeded in a FN-coated μ-Slide VI 0.4 (80606, Ibidi).

Polymorphonuclear cells, consisting mainly of neutrophils, were isolated from whole blood from healthy donors (Sanquin), stored O/N at RT. Briefly, blood was diluted 1:1 in PBS with 5% (v/v) trisodium citrate and pipetted on top of 12.5 mL Percoll (1.047 g/mL) at RT. Cells were centrifuged at 800 g for 20 min (slow start, low brake) and all fractions except neutrophils and erythrocytes were removed. Erythrocytes were lysed twice in ice-cold isotonic lysis buffer (155 mM NH_4_Cl, 10 mM KHCO_3_, 0.1 mM EDTA). Remaining neutrophils were washed with PBS, followed by centrifugation at 450 g for 5 min. Neutrophils were resuspended in HEPES medium (20 mM HEPES, 132 mM NaCl, 6 mM KCl, 1 mM MgSO_4_, 1.2 mM K_2_HPO_4_, 1 mM CaCl_2_, 0.1% D-glucose, 0.5% Albuman (Sanquin Reagents)) and kept at RT for 4h maximum. Neutrophils were prior to use labeled with 2 μM Calcein Red-Orange dye for 20 min at 37 °C. Dye was removed by centrifugation and the neutrophils were directly used. Neutrophils were isolated from four donors.

Two days post-transfection, μ-Slides containing HUVECs were stimulated with 10 ng/ml TNF-α. The slides were connected after 4h to a closed perfusion system to mimic the blood flow. Cells were exposed to flow rates of 0.8 dynes/cm^2^ (HEPES buffer) and kept under 5% humidified CO_2_ atmosphere at 37 °C. Neutrophils where injected in the closed system to allow TEM. 100 μM histamine or a mix of 10 μg/ml ionomycin and 5 mM calcium was used as positive control for the sensor. Fluorescence lifetime was measured using the previously described LIFA setup, with a 40x (Plan Apo, NA 0.95 air) objective. Lifetime stacks in the cyan channel was recorded every 13.5 s, alternated with an image in the red channel. For red excitation a 532 nm LED was used, combined with a 534/20 nm excitation filter, a 575 nm dichroic mirror and a 609/54 nm band-pass filter.

A background correction for removal of background lifetime was done on the lifetime stacks using a manually indicated background region. The background corrected data were used to calculate for each pixel the polar coordinates *M* and *φ*that represent the phase and modulation lifetimes^63^. The polar coordinates were corrected for daily variance compared to the *in vivo* calibration. To this end, the position of the recorded high lifetime state of the positive controls was forced to the position of the high lifetime state of the *in vivo* calibration. The corresponding correction factors (addition for *‘Φ’* and division for *‘M’*) were used to correct all recorded TEM data. Line fraction *‘a’* was calculated and converted into the calcium concentration using the *in vivo* calibration.

Neutrophils were manually tracked in ImageJ (version 1.52k) to collect their intensities. The calcium concentration in the underlying HUVEC was extracted, as well as the cyan intensity. The stage of migration was manually assigned (crawling, diapedesis or post-diapedesis). Since diapedesis is a relatively rapid process, this yields a relatively low number of datapoints. We also measured the concentration before adhesion at the position where a neutrophil would adhere at a later timepoint and assigned the stage ‘before’.

### Organoid imaging

The study was approved by the UMC Utrecht (Utrecht, The Netherlands) ethical committee and was in accordance with the Declaration of Helsinki and according to Dutch law. This study is compliant with all relevant ethical regulations regarding research involving human participants.

Human intestinal organoids from the human ileum (N39; https://huborganoids.nl) stably expressing NLS-mMaroon and nuclear-targeted Tq-Ca-FLITS upon doxycycline induction were generated by lentiviral transduction and maintained as described elsewhere^64^. For differentiation towards EECs, organoids were treated with 1 μg/mL doxycycline for 48 hours, 5 days after seeding. 4 days later (day 6 of differentiation) organoids were analyzed. 10 μg/ml GPBAR-A (4478, Bio-Techne, Abingdon, United Kingdom) was added to the organoids while recording the fluorescence lifetime, using the previously described LIFA setup, with a 40x (Plan Apo, NA 0.95 air) objective. Lifetime stacks were recorded every 10 s. For positive control we added 0.25% (w/v) Triton-X100 together with 12.5 mM CaCl_2_.

Datasets were used to calculate for each pixel the polar coordinates *M* and *φ*that represent the phase and modulation lifetimes^63^. The polar coordinates were corrected for daily variance compared to the *in vivo* calibration. To this end, the position of the recorded high lifetime state of the positive controls was forced to the position of the high lifetime state of the *in vivo* calibration. The corresponding correction factors (addition for *‘Φ’* and division for *‘M’*) were used to correct the data. Line fraction *‘a’* was calculated and converted into the calcium concentration using the *in vivo* calibration.

### Reporting Summary

Further information on research design is available in the Nature Research Reporting Summary linked to this article.

## Data availability

The data produced in this study are available within the article and its Supplementary Information.

All raw data will be available at Zenodo.org upon publication.

Plasmids are deposited for distribution through Addgene (www.addgene.org). The plasmids and corresponding addgene numbers are: pFHL-Tq-Ca-FLITS: #129628, 3xnls-Tq-Ca-FLITS: #129626, Lck-Tq-Ca-FLITS: #129627, pPB-3xnls-Tq-Ca-FLITS: #145030, pLV-H2B-Maroon-P2A-3xnls-Tq-Ca-FLITS: #145027, pInducer-mMaroon-NLS-P2A-3xnls-Tq-Ca-FLITS: being processed.

## Code availability

Custom code and scripts are available through GitHub: https://github.com/Franka-van-der-Linden/Quantitative-Calcium-Imaging.

## Author contributions

F.H.L. and J.G. conceptualized the project, designed the experiments, interpreted the results and wrote the manuscript.

F.H.L., E.K.M., J.A. and J.v.B. participated in the experiments on endothelial cells and were involved in isolation of leukocytes.

J.P, J.D.B, B.P. and H.C. transduced and prepared organoids and assisted with experiments on organoids that were carried out by F.H.L.

A.O.C. generated stable HeLa cell lines.

S.M.A.M. performed FACS.

F.H.L., T.W.J.G. and M.P. were involved in FLIM experiments, performed data analysis and assisted with interpretation of the data.

All authors approved the final manuscript.

## Supplemental Notes

**Supplementary note 1. Construction of the dual expression plasmid pFHL**. To allow fast and easy screening of sensor candidates, a dedicated expression vector was designed, termed pFHL, inspired by the plasmids pDuEx (pDress^35^ were the mTurquoise2, large spatial linker and P2A sequences were removed from the plasmid using NheI restriction sites) and pTorPE^6^. The protein of interest, in this case a candidate sensor, is under control of a CMV promotor for mammalian expression and a rhamnose promotor for bacterial expression. This will eliminate the need of transferring the candidate sensor to a vector for mammalian expression after bacterial screening. At the N-terminus, the sensor is fused to a TorA tag, a 6xHis-tag and an Xpress-tag. The TorA tag primes transport of the sensor to the periplasm of bacteria^65^. As a result, changing the outer environment, for example adding a compound to an agar plate or to a liquid culture, will directly influence the candidate sensor. Also, easy periplasmic isolation will yield a relatively clean protein, ready for quick testing^6^. The 6xHis-tag and the Xpress-tag can be used for protein isolation.

The optimal concentration of rhamnose for expression in *E. coli* using the pFHL vector was determined to be 0.4% (w/v) (**Figure S4A**). The performance of the dual expression vector was verified in *E. coli* and HeLa cells, and compared to the pTorPE plasmid. When expressed in *E. coli*, the R-GECO1 sensor reacted to changing calcium concentration in liquid growth medium (**Figure S4B**). A bigger response was obtained when the sensor was first isolated by isolation of the periplasmid fluid with an osmotic shock. The contrast was lower using the pFHL vector compared to the TorPE vector. However, we found that the contrast was sufficient for screening. The vector also allowed expression in HeLa cells, where addition of ionomycin and extra calcium resulted in a robust intracellular calcium increase, necessary for lifetime measurements (**Figure 1B**).

**Supplementary note 2. Influence of residue 150 in mTurquoise2 on fluorescence lifetime**.

Amino acid V150 is positioned close to the chromophore in mTurquoise2 and therefore we suspected it to affect the fluorescent lifetime of the protein. We randomly mutated this position. Fluorescent and dark colonies were picked and collected on two plates, from which the modulation lifetime was measured using frequency domain FLIM (**Figure S6**) using a custom build FLIM setup and analysis as described before^61^. Lifetimes between 2.4-4.0 ns were recorded among the fluorescent colonies.

**Supplementary note 3. Changing the calcium concentration in the periplasm of bacteria on agar plates**.

*E. coli* cells expressing Tq-Ca-FLITS.0 were grown on LB-agar plates. The modulation lifetime (τ_M_) of the sensor was measured before and > 5 min after 1, 2 or 3 stimulations with a droplet of calcium or chelator EDTA **(Figure S7)**. The lifetime was recorded of the whole plate using a custom build FLIM setup and analysis as described before^61^.

Addition of calcium increased the τ_M_ from 2.70±0.03 ns to 2.83±0.02 ns (mean±sd). Adding more calcium to a colony did not further increase the lifetime. Addition of 200 mM EDTA decreased the lifetime to 2.51±0.07 ns, 2.33±0.08 ns and 2.15±0.11 ns for 1, 2 and 3 drops respectively. The results show that the sensors in bacteria on LB-agar are primarily in the high lifetime state and that the sensors can indeed be influenced from the outer environment, as a result of expression in the periplasmic space.

**Supplementary note 4. Calcium concentrations during transendothelial migration**

Several reports have documented the use of calcium sensitive probes to study changes in calcium levels in endothelial cells upon their interaction with white blood cells. These studies are summarized in **Table S6**.

**Table S6.** Summary of studies that investigate calcium during TEM. The increase in calcium concentration that is observed in endothelial cells varies. This can be partially attributed to the different experimental conditions that are used. Here, we have tried to approach the physiological situation as close as possible by (i) studying TEM under flow at 37 °C, (ii) pretreating the endothelial monolayer with TNF to mimic inflamed conditions, (iii) activating the freshly isolated leukocytes by 20 min incubation at 37 °C and (iv) omitting any of the perturbations that are necessary for labeling cells with fluorescent dyes. Specifically, the use of a genetically encoded probe does not require pre-incubation at room temperature, does not need helper reagents (Pluronic) and omits issues with dye leakage and incomplete hydrolysis. Moreover, Tq-Ca-FLITS uses visible light for excitation (in contrast to Indo-1 and Fura-2 which require UV) and enables direct, intensity-independent quantification. Finally, we have fluorescently labeled both cell types and can therefore precisely analyze the interaction between two cell types and distinguish the different phases of TEM.

| Calcium increase | Endothelial cells | White blood cell | Probe (loading conditions) | Single cell | Flow | Stage | Ref. |
| --- | --- | --- | --- | --- | --- | --- | --- |
| > 800 nM | HUVEC | PMN (10:1) +fMLP | Fura-2 (18 °C & Pluronic) | yes | no | Adhesion | [66] |
| Yes, qualitative | HAEC | PMN (5-10:1) | Indo-1 (23 °C, RT) | no | no | Adhesion | [67] |
| Yes, qualitative | HUVEC | CD65+/NK (20:1 ratio) | Fluo-3 (22 °C & Pluronic) | yes | no | Adhesion | [68] |
| 180 nM | Lung HMVEC | Lymphocyte (10<sup>5</sup> /ml) | Fura-2 (37 °C & Pluronic) | no | no | Adhesion | [69] |
| Qualitative: ~60% of cells | HMEC-1 | Monocytes (10:1) | Indo-1 (RT) | yes | no | Adhesion | [70] |
| ~200 nM | HAEC | Monocytes | Fura-2 (24 °C) | yes | no | Adhesion | [71] |
| Qualitative: ~25% of cells | HUVEC | Neutrophils | YCaM | yes | yes | Rolling | [72] |
| Qualitative: None of the cells | HUVEC | Neutrophils | YCaM | yes | yes | Crawling | [72] |
| No: < 80 nM | HUVEC | Neutrophils | Tq-Ca-FLITS | yes | yes | Crawling & diapedesis | This study |

## Supplemental Movies

**Supplementary movie S1:** Calcium calibration in HeLa cells that express a nuclear tagged Tq-Ca-FLITS biosensor. The intensity and phase lifetime data are shown after cell permeabilization and equilibration at different external calcium concentrations.

**Supplementary movie S2:** Calcium levels in endothelial cells monitored with plasma membrane targeted Tq-Ca-FLITS before and after stimulation with 1 µM histamine. Displayed are on the left the intensity of the probe is shown and on the right the calcium concentration calculated from the lifetime data in false color according to the color scale.

**Supplementary movie S3:** Calcium levels in endothelial cells monitored with plasma membrane targeted Tq-Ca-FLITS during transendothelial migration. Displayed are on the left the intensities of the probe and the neutrophil (in cyan and red respectively) and on the right the calcium concentration calculated from the lifetime data in false color according to the color scale. The white region of interest shows the location of the neutrophil, showing that calcium is not elevated in this region.

**Supplementary movie S4:** Calcium changes measured in nuclei of human small intestinal organoids stimulated with 10 μg/ml GPBAR-A at 25 sec. Displayed are on the left the intensity of the probe and on the right the calcium concentration calculated from the lifetime data in false color according to the color scale.

